# Covalent inhibition by a natural product-inspired latent electrophile

**DOI:** 10.1101/2023.01.16.524242

**Authors:** David P. Byun, Jennifer Ritchie, Ronald Holewinski, Hong-Rae Kim, Ravichandra Tagirasa, Joseph Ivanic, Claire M. Weekley, Michael W. Parker, Thorkell Adresson, Euna Yoo

## Abstract

Strategies to target specific protein cysteines are critical to covalent probe and drug discovery. 3-bromo-4,5-dihydroxazole is a natural product-inspired, synthetically accessible electrophilic moiety that has previously been shown to react with nucleophilic cysteines in the active site of purified enzymes. Here we define the global cysteine reactivity and selectivity of a set of 3-bromo-4,5-dihydroxazole-functionalized chemical fragments using competitive chemoproteomic profiling methods. Our study demonstrates that 3-bromo-4,5-dihydroxazoles capably engage reactive cysteine residues in the human proteome and the selectivity landscape of cysteines liganded by 3-bromo-4,5-dihydroxazoles is distinct from that of haloacetamide electrophiles. Given its tempered reactivity, 3-bromo-4,5-dihydroxazoles showed restricted, selective engagement with proteins driven by interactions between a tunable binding element and the complementary protein sites. We further validate that 3-bromo-4,5-dihydroxazoles form covalent conjugates with glutathione *S*-transferase Pi (GSTP1) and peptidyl-prolyl *cis*-*trans* isomerase NIMA-interacting 1 (PIN1), emerging anti-cancer targets. Together, this study expands the spectrum of optimizable chemical tools for covalent ligand discovery and highlights the utility of 3-bromo-4,5-dihydroxazole as a cysteine-reactive electrophile.

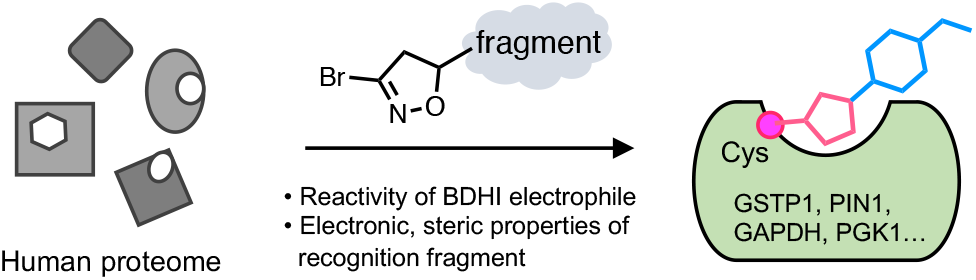

## INTRODUCTION

Covalent ligands that combine chemical reactivity and molecular recognition present a powerful strategy for chemical probes and drug development.^1^ Cysteine-directed covalent warheads are already a component of many FDA-approved drugs and advanced chemoproteomic studies have uncovered thousands of additional cysteines that can be covalently modified by small molecules.^2-5^ However, these sites have been identified using a relatively limited number of electrophilic chemotypes, most notably haloacetamides, nitriles, and acrylamides.^6^ Thus, there remains an unmet need for novel electrophilic scaffolds whose reactivity and selectivity can be rationally tuned to address protein targets of interest.

Electrophilic natural products have proven a remarkable source of chemical innovation and anti-pathogenic/-tumorigenic activity.^7, 8^ One unique naturally-occurring electrophile is the 3-chloro-4,5-dihydroisoxazole heterocycle, a five-membered ring found in the natural product acivicin. Acivicin was originally isolated from the fermentation broth of *Streptomyces sviceus*, and exhibits anti-inflammatory, anti-bacterial, anti-parasitic, and anti-cancer activities.^9, 10^ Mechanistic studies have found that the 3-halo-4,5-dihydroisoxazole scaffold of acivicin can react with cysteine residues in enzymes such as glutamine amidotransferase to form a covalent bond by displacement of the chlorine atom (Figure 1A).^11-16^ It has been hypothesized that protein binding-associated activation and stabilization precede the covalent modification of acivicin’s targets and confer specificity to this heterocyclic warhead.^11, 17^ However, besides a single effort to identify direct targets of acivicin itself,^18^ the potential of the 3-halo-4,5-dihydroisoxazole scaffold to globally address unique targets in the human proteome has not been assessed.

**Figure 1.**
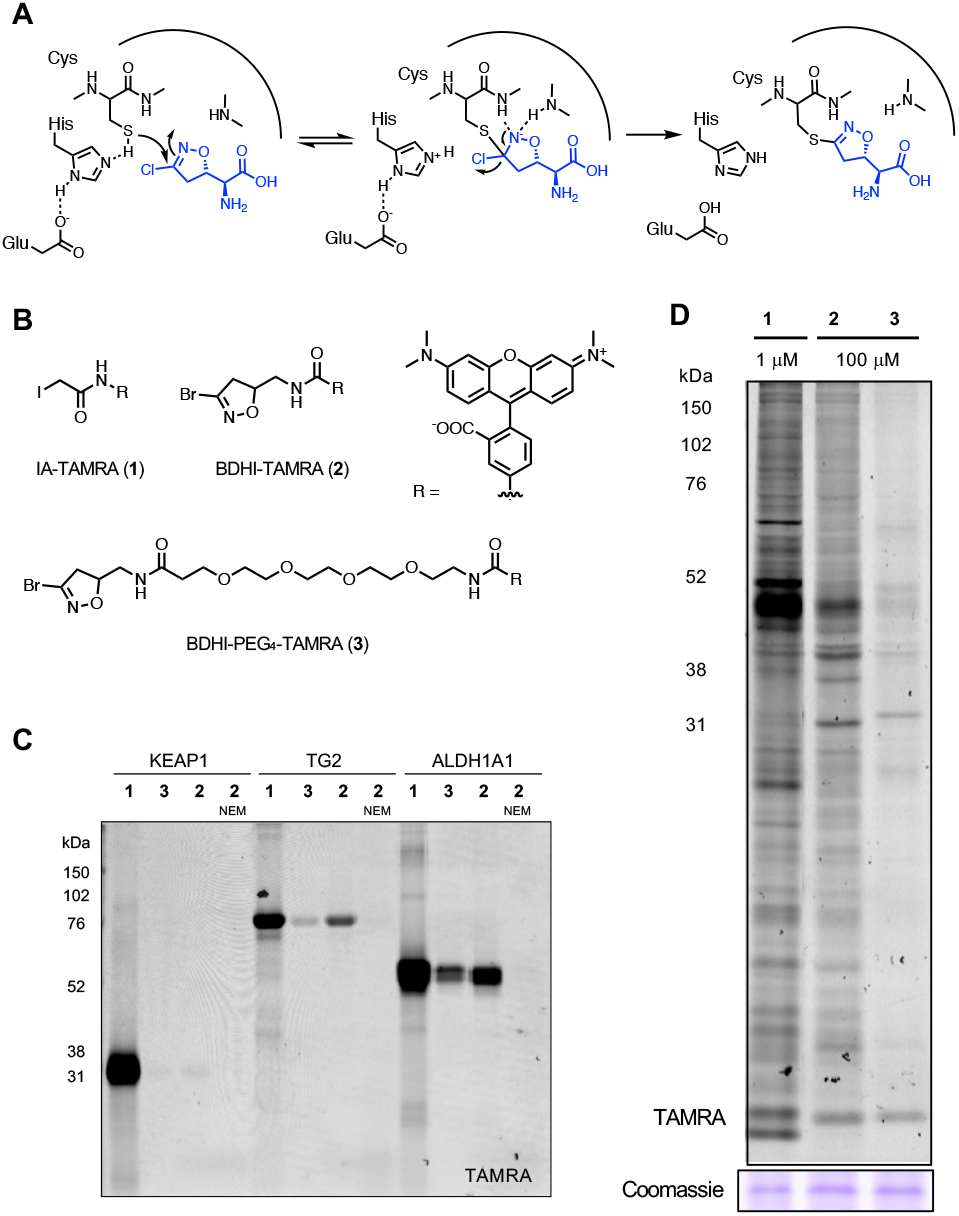
BDHI electrophile reacts with proteins in the human proteome.(A) Covalent inactivation mechanism of acivicin for glutamine amidotransferase. (B) Structure of fluorescent-BDHI probes. (C) In-gel fluorescence scanning of recombinant KEAP1, TG2 and ALDH1A1 labeled with BDHI-TAMRA (**2**) and BDHI-PEG_4_-TAMRA (**3**) probes (100 μM, 2 h). (D) In-gel fluorescence image depicting protein bands labeled with fluorescent-BDHI probes in the human proteome. Soluble proteome from Jurkat cells was treated with the indicated probes for 2 h, followed by SDS-PAGE and in-gel fluorescence scanning.

Here we set out to define and manipulate the reactivity and selectivity of this natural product-inspired electrophilic scaffold using a combined rational design and proteome-wide covalent ligand screening approach. First, we explore the attenuated reactivity of a 3-bromo-4,5-dihydroisoxazole (BDHI) scaffold towards human proteins. Analysis of this electrophile’s quantum chemical properties enables the design of chemoproteomic probes with orthogonal glutathione reactivity, solution stability, and proteomic reactivity. Applying this knowledge, we synthesize a small panel of BDHI analogs and analyze their properties using competitive chemoproteomic profiling approach. This reveals the ability of selected fragments to drive site-specific engagement of a distinct subset of cysteines in the human proteome, whose functional significance could be validated via biochemical assays and structural modeling. Overall, our studies demonstrate the potential for natural product-inspired electrophiles to empower inverse drug design strategies and provide a foundation for further applying the BDHI scaffold to advance ligand discovery efforts.

## RESULTS AND DISCUSSION

### Selective cysteine reactivity of the BDHI electrophile

We selected the 3-bromo-4,5-dihydroisoxazole (BDHI) moiety as a subject for our studies because it is slightly more reactive than the chloride-substituted counterpart but is not expected to alter target preferences.^19^ To benchmark the reactivity of the BDHI electrophile, we compared a general fluorescent cysteine labeling reagent (**1**) to two fluorescent BDHI probes (**2** and **3**; Figure 1B). As recombinant protein targets we chose Kelch-like ECH-associated protein 1 (KEAP1), a protein that forms adducts with many electrophilic small molecules and contains 24 cysteine residues, as well as transglutaminase 2 (TG2) and aldehyde dehydrogenase 1A1 (ALDH1A1), two enzymes that have been previously reported to interact with the BDHI electrophile which contain 20 and 11 cysteine residues, respectively.^13, 19^ Proteins were individually incubated with 100 μM **1**-**3** for 2 h at 37°C and analyzed by SDS-polyacrylamide gel electrophoresis (SDS-PAGE). In-gel fluorescence scanning indicated that iodoacetamide (IA) probe **1** capably reacts with all three proteins, whereas BDHI probes **2** and **3** only demonstrate visible labeling of TG2 and ALDH1A (Figure 1C). The hydrophobic BDHI-tetramethylrhodamine (TAMRA) analogue **2** exhibited more intense protein labeling than BDHI-PEG_4_-TAMRA **3**. Of note, *N*-ethylmaleimide (NEM) pretreatment completely abolished the protein labeling by probe **2**, consistent with cysteine-directed labeling. When incubated with proteomic lysates of Jurkat cells (a human T cell lymphoma line), BDHI-TAMRA probes **2** and **3** formed conjugates with numerous proteins, with a few showing preferential engagement relative to IA-TAMRA **1** (Figure 1D). BDHI probes labeled proteins to a much lesser extent compared to promiscuous iodoacetamide probe and required a relatively higher concentration, likely due to low or slow reactivity of the BDHI electrophile towards target proteins with a non-optimal binding element. It is also possible that multiple cysteines present in each protein are modified with a more reactive IA electrophile, whereas the BDHI electrophile only reacts more selectively with cysteine(s) in a defined binding pocket. These studies provide the first human proteome-wide characterization of the natural product-inspired BDHI scaffold and led us to further explore its tunable selectivity.

### Computational analysis guides tuning of BDHI reactivity

The five-membered heterocyclic ring of BDHI is unique relative to smaller electrophiles (e.g. haloacetamides) in that it is of similar size to a pharmacophore and has the potential to contribute to molecular recognition of targets itself. Acivicin inhibits glutamine amidotransferase by placing a 3-chloro-4,5-dihydroisoxazole directly adjacent to an amino acid backbone, which can influence both the electrophilicity and molecular recognition of protein surfaces by its covalent warhead. Considering this, we hypothesized that computationally-informed structure-function analysis may be able to similarly inform the interplay of these properties in the BDHI electrophile. For these studies, we designed two structurally distinct BDHI probes, **4** and **5** (Figure 2A). Compound **4** contains an alpha amido-BDHI which is structurally identical to the warhead used in our fluorescent probes. Compound **5** replaces the amide with a substituted phenyl ring, which confers greater lipophilicity and electron-withdrawing properties. Each compound was also designed to incorporate a bioorthogonal alkyne handle to enable experimental analyses of protein labeling. Density functional theory (DFT) calculations were used to predict the susceptibility of these compounds towards nucleophilic attack by thiols and understand possible changes in reactivity resulting from differential substitution. Analyzing the reaction energy profiles of each compound with a simple methanethiol, we found that there is large transition state barrier (45.5 kcal/mol for compound **4** and 44.3 kcal/mol for compound **5**) to react with MeSH (Figure 2B). This is in good agreement with the observation that BDHIs do not readily react with glutathione (GSH) at pH 7.4 where the equilibrium lies on the thiol side given the p*K*a of GSH (9.17).^20^ On the other hand, nucleophilic addition of the thiolate anion (MeS^-^) is feasible and likely spontaneous (ΔG^‡^ = 4.4 and 7.7 kcal/mol for **4** and **5**, respectively; Figure 2C), implicating thiol deprotonation as the likely rate-limiting step. Given this, we speculate that BDHIs may preferentially react with hyper-reactive cysteines with reduced p*K*a (heightened nucleophilicity).^21^ Overall energy profiles are very similar between **4** and **5**; however, **5** has slightly stronger binding energy for initial reactant complex with MeS^-^ (by 1.8 kcal/mol).

**Figure 2.**
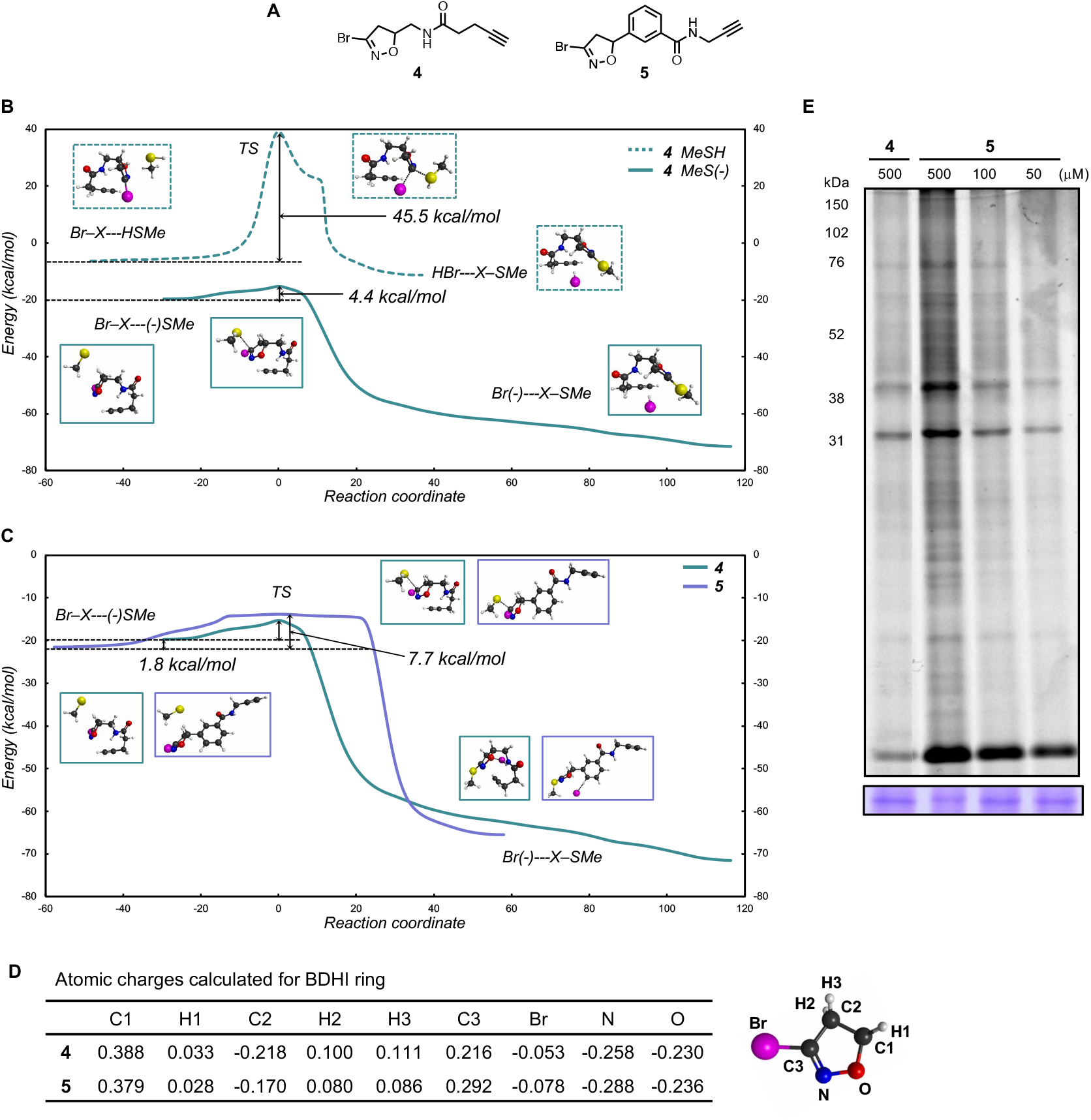
BDHI-alkyne probes. (A) Structures of BDHI-alkyne probes **4** and **5**. (B-C) Computed reaction energy profile of the BDHI electrophile with thiol nucleophiles using DFT calculations. Comparison of the reactions between MeSH and MeS^-^ with **4** (B) and comparison of the reactions between **4** and **5** with MeS^-^ (C). (D) Charge analysis of indicated atoms in the BDHI ring of compound **4** and **5**. (E) In-gel fluorescence image depicting protein bands labeled with BDHI-alkyne probes in the human proteome. Soluble proteome from Jurkat cells was treated with the indicated probes for 2 h and subsequently subjected to CuAAC with a TAMRA-N_3_, followed by SDS-PAGE and in-gel fluorescence scanning.

Next, we calculated the atomic charges of BDHI ring of the two probes by fitting to the electrostatic potentials. Importantly, N-O bond of the dihydroisoxazole (DHI) ring is highly electronegative and contributes to the polarization of the C3-C2 bond, and the C3 imino carbon features a positive electrostatic profile (numbering shown in Figure 2D). The electronegative potential of DHI ring may potentiate binding to oppositely charged protein pockets and ultimately allow the stabilization of a tetrahedral intermediate formed upon nucleophilic attack. Comparison of **4** and **5** observed an overall similar electrostatic potential, whereas an obvious difference was detected at the C3 carbon. Compound **5** exhibited a higher electropositive potential at the C3 compared to **4** suggesting slightly heightened susceptibility to nucleophilic attack. This is consistent with our analysis of the MeS^-^/BDHI reaction coordinate. To experimentally assess the proteome reactivity of these two probes, we synthesized each and treated Jurkat proteomes with BDHI alkyne probes **4** and **5** followed by conjugation with a fluorescent tag using a copper-catalyzed azide-alkyne cycloaddition (CuAAC) chemistry. Each compound displayed efficient labeling of multiple proteins (Figure 2E). However, consistent with its predicted heightened reactivity and lipophilicity, probe **5** showed a broader labeling profile than **4**. These studies demonstrate a computationally-guided strategy for manipulating the reactivity of a natural product-inspired electrophile and provide access to a set of reporters for assessing the protein reactivity of two orthogonal BDHI chemotypes.

### Proteome reactivity of BDHI-containing fragments

Based on our analysis of fluorescent and biorthogonal BDHI probes, we envisioned that BDHI warheads could be used to produce small molecules capable of selectively reacting with distinct subsets of the human proteome. To explore this notion, we installed the BDHI electrophile onto a set of chemical fragments (so called “scout fragments”) previously shown to drive covalent protein engagement with different electronic and steric properties^2^ and assessed their proteomic reactivity. BDHI fragments were synthesized using 1,3-dipolar cycloaddition reactions with a stable precursor dibromo-formaldoxime to afford a racemic mixture of 5-substituted 3-bromo-isoxazole derivatives (Scheme S1 and Figure 3A).^22, 23^ This synthetic procedure also allows to directly convert acrylamide-bearing covalent ligands to corresponding BDHI analogs (**6**-**10**). Analysis of the electrostatic potential surface of this small panel of BDHI analogs again identified slight differences in the electrostatic potential profile at the BDHI ring, with more substantial differences at the C1 carbon as well as at the substituted aromatic moieties (Figure S1 and Table S1). This suggests 5-substitution may contribute not only to the steric effect (by altering how binding moieties present the BDHI electrophile) but also to the electronic effect for binding.

**Figure 3.**
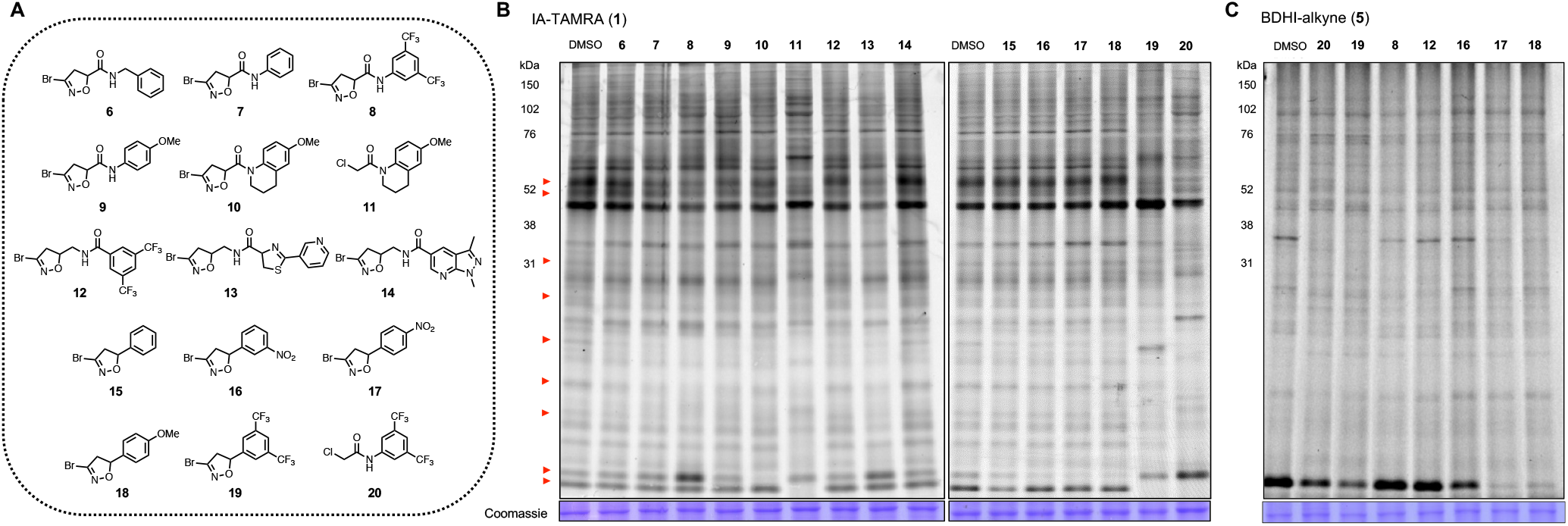
BDHI electrophile-containing fragments competitively block the labeling of proteins. (A) Structure of BDHI-functionalized fragments. (B) Initial competitive analysis of the proteomic reactivity of fragments using an IA-TAMRA. Soluble proteome from Jurkat cells was treated with the indicated fragments (500 μM each) for 4 h, followed by labelling with IA-TAMRA (1 μM, 1 h) and analysis by SDS-PAGE and in-gel fluorescence scanning. Red arrows highlight protein bands that showed diminished IA-TAMRA labeling. (C) BDHI ligands also competitively block the labeling of proteins by BDHI-alkyne **5** (100 μM, 2 h, subsequently conjugated with N_3_-TAMRA).

To quickly interrogate the cysteine reactivity of these compounds, we used a competitive gel-based screening, incubating Jurkat cell lysates with each compound followed by labeling of cysteine with IA-TAMRA. In this approach, compound pretreatment that competitively blocks probe labeling results in reduced fluorescent labeling of individual proteins. Given their low molecular weight, we screened them at a single high concentration (500 μM) similar to those used in fragment-based ligand discovery.^24^ Visual inspection of protein bands (Figure 3B, red arrows, left to right) revealed a discrete subset of proteins whose IA-TAMRA labeling was diminished in intensity upon pre-treatment of BDHI analogs. However, compared to α -chloroacetamide (CA)-containing compounds **11** (KB02) or **20** (KB03),^2^ BDHI analogs exhibited more restricted and selective blockade of IA-TAMRA-protein interactions. Overall, IA-TAMRA labeling was altered most substantially by pretreatment with **8** and **13**. In contrast, compound **14** did not show any visible evidence of competition, demonstrating the high dependence of BDHI reactivity on cognate fragment scaffold. When compared to fragments that contain a mildly reactive acrylamide electrophile (**21**-**23**; Figure S2), slightly enhanced proteomic reactivity was observed from the BDHI-containing fragments. Blockade of IA-TAMRA-protein interactions by 5-phenyl derivatives (**15**-**19**) was overall less pronounced. Considering the possibility that this reflects differential labeling kinetics of IA-TAMRA versus BDHI electrophiles, we also assessed competitive labeling of proteins by BDHI-TAMRA probe **2** (Figure S3) and alkyne probe **5** (Figure 3C). This experiment confirmed competitive labeling of many proteins by 5-phenyl analogs, with **18** showing the broadest level of proteome-wide reactivity. Additional alkyne-functionalized analogs of selected fragments (**24**-**26**) were synthesized and directly assessed for their ability to label proteins (Figure S4). Reflecting the competition data, alkynylated analog of **7** (**25**) showed efficient labeling of many proteins with enhanced reactivity. Supporting the attenuated reactivity of BDHI electrophile towards sulfhydryl groups in solution and a requirement for molecular recognition in BDHI labeling, we found that incubating three BDHI analogs (**8, 10**, and **18**) with GSH at pH 7.4 did not show any formation of a covalent adduct, whereas CA-containing compounds **11** and **20** produced more than 50% of corresponding adducts within 4 h of incubation (Figure S5A). To our surprise, BDHI **19** with a bis(trifluoromethyl)phenyl group was found to be unstable in phosphate buffer (Figure S5B). As such, we excluded compound **19** from further studies. Together, these data suggest that BDHI is a mildly reactive electrophile that can acts as a covalent protein modifier upon interaction within protein binding pockets and its proteomic reactivity and selectivity can be largely modulated by fragments appended to the warhead.

### Competitive reactivity profiling defines a subset of proteomic cysteines targeted by BDHI analogs

Based on our observation on competitive labeling using IA-TAMRA, we next performed mass spectrometry (MS)-based chemoproteomic profiling to globally map the protein targets of our BDHI probes. In brief, cell lysates were treated with either vehicle (DMSO) or a BDHI-containing ligand (Figure 4A). Samples were then labeled with a cysteine reactive IA-desthiobiotin (DTB) probe,^4^ followed by digestion and tandem mass tags (TMT) labeling. After combining samples, labeled peptides were enriched by streptavidin beads, eluted, and subjected to LC-MS/MS analysis. Cysteines that show reduction in TMT signals for BDHI-treated samples compared to DMSO control (calculated as a competition ratio) are considered candidate covalent targets. We selected IA-DTB as the chemoproteomic probe (as opposed to **4** or **5**) due to its broad validation and to avoid potential technical issues resulting from poor ionization and fragmentation of the BDHI-adduct.^4^ Based on gel-based data, we selected compounds **8, 10, 13, 14**, and **18** for analysis and compared their reactivity to promiscuous control compounds KB02 (**11**) and KB03 (**20**). Experiments were carried out by incubating compounds (500 μM, 4 h, in triplicate) in Jurkat whole cell lysates. Cysteine occupancy was determined by log_2_ fold change in the abundance of TMT reporter ion signals (Table S2). Using this approach, we identified 237, 35, 9, 0, and 17 liganded cysteine sites that showed > 75% reduction in TMT signals (log_2_ FC > 2, *p* value < 0.05) after incubation with BDHI **8, 10, 13, 14**, and **18**, respectively (Figures 4B and S6A). This number is significantly less than that observed for haloacetamide **11** (2209 sites) or **20** (3306 sites), again reflecting the more restricted blockade of IA probe-protein interactions by BDHI derivatives. Consistent with gel-based analyses, IA-DTB profiling revealed **8** and **14** to be the most and least reactive analogs, respectively. BDHIs were found to target functional proteins including both enzymes and non-enzymes. A large fraction of non-catalytic proteins we identified were proteins that are involved in mediating protein-protein or protein-nucleic acid interactions. We found 21 BDHI-liganded sites among the top 150 most reactive cysteine residues identified from previous study,^25^ including the active site cysteines of glutathione *S*-transferase omega-1 (GSTO1) and methylated DNA protein cysteine methyltransferase (MGMT) (Figure S6B). Moreover, compounds were observed to engage discrete sets of both overlapping and distinct cysteines (Figures 4C-D and S6C-D). A brief survey of these targets follows:

**Figure 4.**
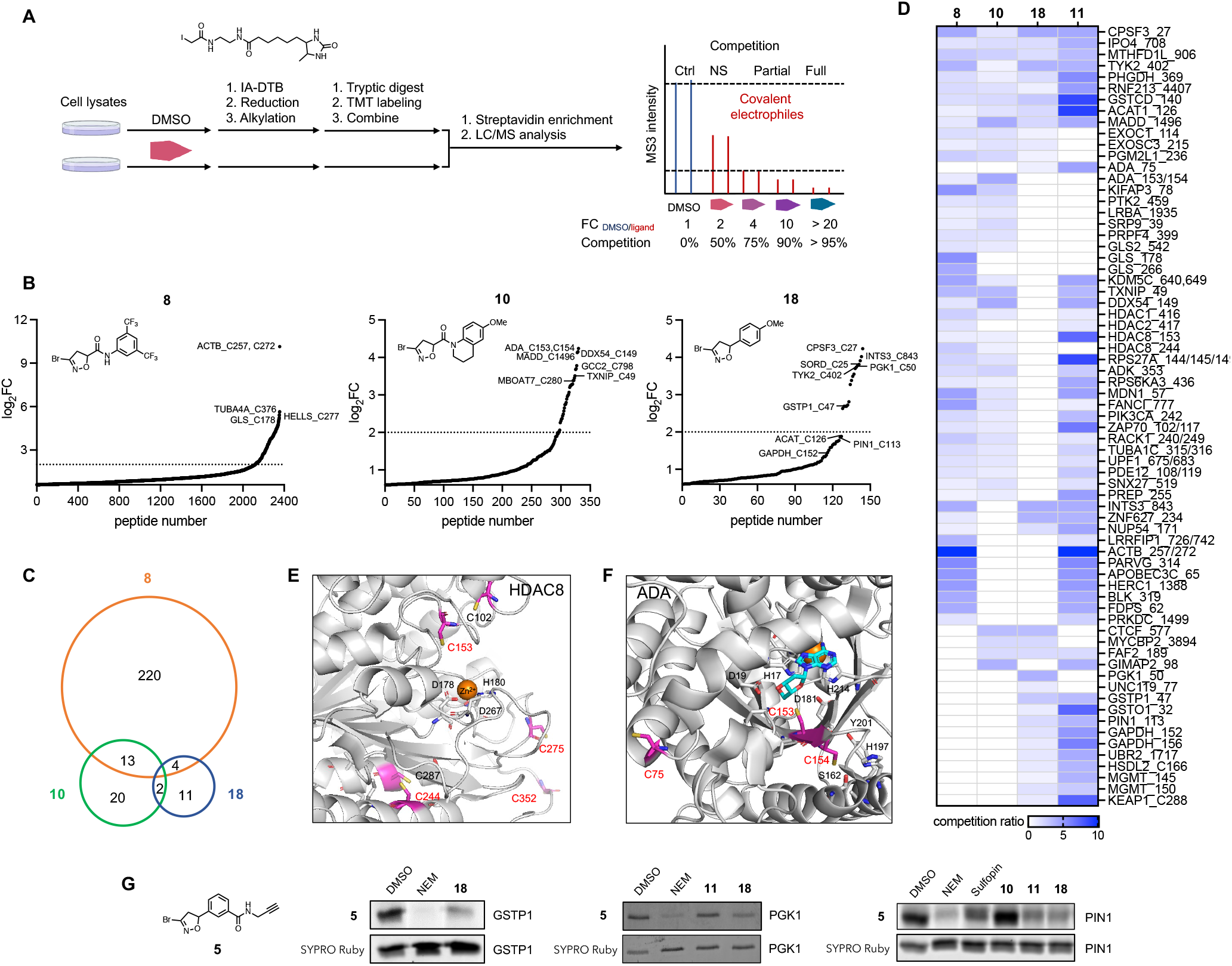
Competitive MS-based reactivity profiling defines cysteines liganded by BDHI analogs. (A) Workflow for competitive measurement of cysteine ligandability with fragment electrophiles using IA-DTB probe. Steps include treatment of cell lysate with compounds (500 μM, 4 h), treatment with IA-DTB probe (500 μM, 2 h), trypsin digestion, TMT labeling, combination, avidin enrichment for biotinylated cysteine-containing peptides and analysis of competition ratio. (B) Structures and competition ratios (log_2_FC values) from Jurkat cell proteome treated with BDHI **8, 10**, and **18**. (C) Venn diagram representation of number of cysteine sites significantly liganded (log_2_FC > 2) by BDHIs. Results were obtained by comparing the site overlap at a given competition ratio threshold for each ligand. (D) Heatmap of competition ratios for representative cysteines and fragments. (E) Crystal structure of HDAC8 (PDB: 2V5W). The zinc binding domain is shown along with reactive cysteines (depicted in magenta). (F) Crystal structure of substrate bound, human ADA (PDB: 3IAR) with possible cysteine sites for covalent modification. Adenosine is depicted in cyan. (G) Competition binding assay between BDHI **18** and BDHI alkyne **5**. Recombinantly expressed and purified PGK1, GSTP1, PIN1 were treated with each compound (500 μM, 4h), labeled with BDHI **5** (500 μM, 2h) and conjugated with TAMRA-azide for in-gel fluorescence scanning.

- Cysteines that were liganded by both haloacetamide **11** and a subset of BDHI analogs include C27 of cleavage and polyadenylation specificity factor subunit 3 (CPSF3), C708 of importin-4 (IPO4), C906 of mitochondrial monofunctional C1-tetrahydrofolate synthase (MTHFD1L, also known as formyltetrahydrofolate synthetase), and C140 of glutathione *S*-transferase C-terminal domain-containing protein (GSTCD). However, we found that while the CA scout fragments **11** and **20** target multiple cysteines in each protein, BDHI derivatives appear to selectively engage only one cysteine. For example, BDHI **18** showed preferential engagement with the active site C126 among the four reactive cysteines (C119, C126, C196, and C413) found in the mitochondrial acetyl-CoA acetyltransferase (ACAT1, Figure S7A). On the other hand, for glyceraldehyde 3-phosphate dehydrogenase (GAPDH), BDHI **18** engaged with both C152 and C156 residues (Figure S7B).
- Among the liganded sites that were uniquely observed with BDHIs were C178 and C266 of kidney-type glutaminase (GLS) and C244 of histone deacetylase 8 (HDAC8) with **8**, C153/154 of adenosine deaminase (ADA) with **10**, and C50 of phospho-glycerate kinase 1 (PGK1) with **18**. It is noteworthy that acivicin as a glutamine mimetic has been reported to bind to the active site of GLS and other glutamine-binding enzymes.^12, 26, 27^
- HDAC8, which belongs to the human histone deacetylase family consisting of 11 isozymes, is a validated target for T cell lymphoma.^28^ In addition, pharmacological inhibition of HDAC8 has been demonstrated to enhance antitumor immunity and efficacy of immunotherapy.^29^ The enzymatic activity of HDAC8 is regulated by a reversible thiol/disulfide redox switch involving C102 and C153 residues (Figure 4E).^30^ Three additional pairs of cysteines (C125/C131, C244/C287, and C275/C352) are in proximity that could also potentially form further disulfide bonds. C244, which is unique to HDAC8, is one of the most buried cysteines with the highest calculated p*K*_a_ value.^31^ It is thus surprising that this cysteine with theoretically low reactivity showed ligandability with BDHI compound **8**. Interestingly, a screening of maleimide analogs appended with the 3,5-bis(tri-fluoromethyl)phenyl group (similar to **8)** has shown to label C244 and C275 of HDAC8 and inhibit the enzyme activity.^32^ Further validation and investigation will be required to understand the role of C244 in ligand binding and modulation of HDAC8 activity.
- ADA plays an important role in purine metabolism by catalyzing the hydrolytic deamination of adenosine to inosine. It is highly expressed in T lymphocytes and known to regulate T cell co-activation through its interaction with CD26 (DPP4) at the cell surface.^33, 34^ While covalent modification of C75 by electrophilic small molecules has been reported to allosterically inhibit the enzymatic activity of ADA and lead to antiproliferation of lymphocytic cells,^35^ there was no covalent modifier identified to target C153 or C154 that is located close to the active site of ADA (Figure 4F).
- PGK1 is the first ATP-generating enzyme in the glycolytic pathway associated with one-carbon metabolism and cellular redox regulation. PGK1 is involved in shaping the inflamed tumor microenvironment and mediating interaction between tumor metabolism and immunoediting. Among the seven cysteines found in PGK1 (Figure S7C), C50 has been reported to undergo *S*-sulfinylation (-SO_2_H) during oxidative stress.^36^ Under hypoxic condition that triggers endogenous H_2_O_2_ production, PGK1 is translocated into the mitochondria.^37^ Whether the modification of this cysteine has any impact on mediating mitochondrial translocation and/or kinase activity of PGK1 remains to be investigated.

To provide orthogonal validation for target interactions we employed BDHI alkyne probe **5**. When incubated with the recombinant protein followed by a click reaction with TAMRA-azide, compound **5** showed detectible labeling of PGK1, GSTP1, and PIN1, which in each case was blocked by pretreatment with the cognate BDHI fragment **18** (Figure 4G). Overall, these data provide evidence that the BDHI electrophile, when coupled to a suitable binding element, has the potential to site-specifically target cysteines. Our analysis also highlights a small set of cysteines that show exclusive engagement with the BDHI electrophile.

### Functional engagement of proteins by the BDHI electrophile

Next, we sought to define the functional effects of covalent adduction by the BDHI electrophile. For these studies we prioritized targets based on their therapeutic relevance, availability of structures in the Protein Data Bank, and overall abundance which lends further confidence to chemoproteomic identifications. This led us focus on glutathione *S*-transferase Pi (GSTP1) and peptidyl-prolyl *cis*-*trans* isomerase NIMA-interacting 1 (PIN1). GSTP1 is the most prevalent cytosolic glutathione transferases which catalyzes the conjugation of glutathione to electrophilic compounds as a cellular detoxification process.^38^ GSTP1 is also involved in regulating cellular redox state and intracellular signal transduction.^39^ Owing to its overexpression in a variety of malignant cells and confirmed role in promoting tumorigenesis, GSTP1 inhibitors have emerged as promising cancer therapeutic agents.^40-42^ Among the four cysteine residues found in GSTP1, C47 and C101 have each been demonstrated to be solvent accessible, highly reactive, and targeted by irreversible inhibitors (Figure 5A).^43^ Specifically, the chemical modification of C47, a residue located near the active site, has been shown to cause loss of enzyme activity.^44, 45^ We used recombinantly expressed human GSTP1 to confirm that BDHI **18** and the related 5-phenyl-substituted analog **15**, but not **8** or **10**, inhibit the labeling of GSTP1 by IA-TAMRA, consistent with the competition observed in MS-based profiling experiments (Figure 5B). Consequently, we carried out enzyme activity assays and found that BDHI **18** inhibits GSTP1 enzyme activity in vitro with an IC_50_ of 15 μM, determined by monitoring the transfer of GSH to 1-bromo-2,4-dinitrobenzene (BDNB) (Figure 5C). To confirm the site of modification, recombinant GSTP1 was incubated with BDHI **18** at 37°C for 4 h similar to the labelling conditions and subjected to trypsin digestion and LC-MS/MS analysis. All data were searched for a differential modification mass of 175.06 (covalent adduct formation through the Br replacement of BDHI **18**). This led to the identification of two cysteine residues, the active site residue C47 as well as C101, as modified by **18** in GSTP1 (Figures 5D and S8). To further validate and understand this interaction we obtained multiple crystal structures from GSTP1 crystals soaked with compound **18**. While the position of **18** was not unambiguously determinable, structures showed a loss of electron density in the α 2 helix (residues 35-50) and an opening of the interface between the two monomers upon treatment with **18** (Figure S9 and Table S3). The loss of electron density was increased with fragment concentration and soaking time. This observation is consistent with the fragment interacting at the C47 in the α 2 helix and possibly with C101 at the dimer interface. Similar structural changes have been observed in GSTP1 with compounds that bind C47 and/or C101 in the absence of GSH.^46-48^ Mass spectrometric analysis of crystals soaked with **18** alone or back-soaked with GSH indicated that a majority of the protein in the crystals is bound to one or two molecules of **18** (Table S4), again consistent with these effects being driven by ligand-binding.

**Figure 5.**
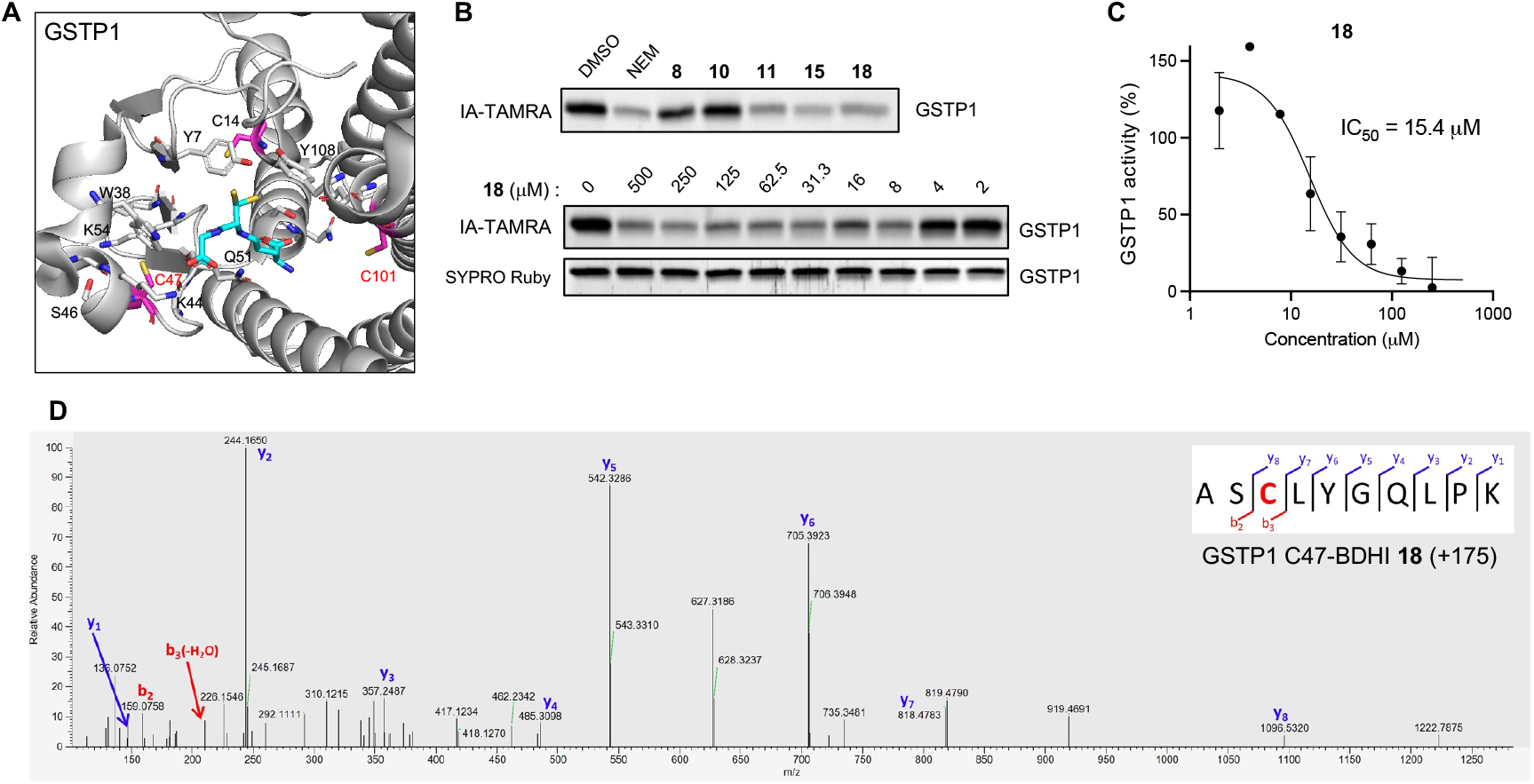
Validation of GSTP1 as the BDHI-reactive protein. (A) Crystal structure of GSTP1 (PDB: 6LLX). Reactive cysteine sites (in magenta) are shown along with binding of glutathione depicted in cyan. (B) Competitive and concentration-dependent inhibition of GSTP1 labeling by **18**. Recombinant GSTP1 was preincubated with each compound (500 μM, 4h), labeled with IA-TAMRA (1 μM, 1 h) and subsequently analyzed by SDS-PAGE and in-gel fluorescence. (C) Inhibition of GSTP1 activity determined by monitoring the transfer of GSH to 1-bromo-2,4-dinitrobenzene. (D) Annotated MS2 fragmentation spectra analysis of GSTP1 liganded with BDHI **18** at Cys47, highlighted in red.

As a second target we analyzed the binding of BDHI **18** to PIN1. PIN1 is a human peptidyl-prolyl *cis*-*trans* isomerase (PPIase) that is overactivated in numerous tumor types and whose aberrant activation has been associated with tumorigenesis.^49, 50^ The PIN1 active site contains a nucleophilic cysteine residue (C113) that can be targeted by covalent inhibitors. These molecules have been shown to exhibit antiproliferative effects in various cancer cell lines including neuroblastoma and pancreatic ductal adenocarcinoma.^51, 52^ Using a gel-based assay with recombinant protein, we found that BDHI **18** inhibits PIN1 labeling by IA-TAMRA in a dose-dependent manner (Figure 6A). To further confirm that BDHI **18** site-specifically engages PIN1, we performed LC-MS/MS analysis and validated the catalytic residue Cys113 as the site of labeling (Figure 6B). In order to better understand the structural basis for PIN1 engagement, we employed *in silico* modeling to predict potential binding mode of BDHI **18** to PIN1. Comparing the lowest energy binding pose of **18** to that of sulfopin,^52^ an inhibitor of PIN1 that contains haloacetamide warhead, we found that the BDHI warhead can covalently engage C113 with binding driven by hydrogen bond interactions of the DHI ring with neighboring S115 (Figure 6C). In this pose, the 4-methoxyphenyl group of **18** occupies the hydrophobic binding pocket surrounded by M130, Q131, T152 and H157 in a manner near identical to the sulfolane ring of sulfopin, while also making a hydrogen bond interaction with the backbone amide of Q131. These data confirm that BDHI electrophile targets and covalently modifies cysteine residues in the allosteric and catalytic site of GSTP1 and PIN1. Although further optimization is needed to improve the potency of these compounds, our study demonstrates that BDHI compounds have the potential for the development of novel covalent inhibitors of GSTP1 and PIN1, emerging anti-can-cer targets.

**Figure 6.**
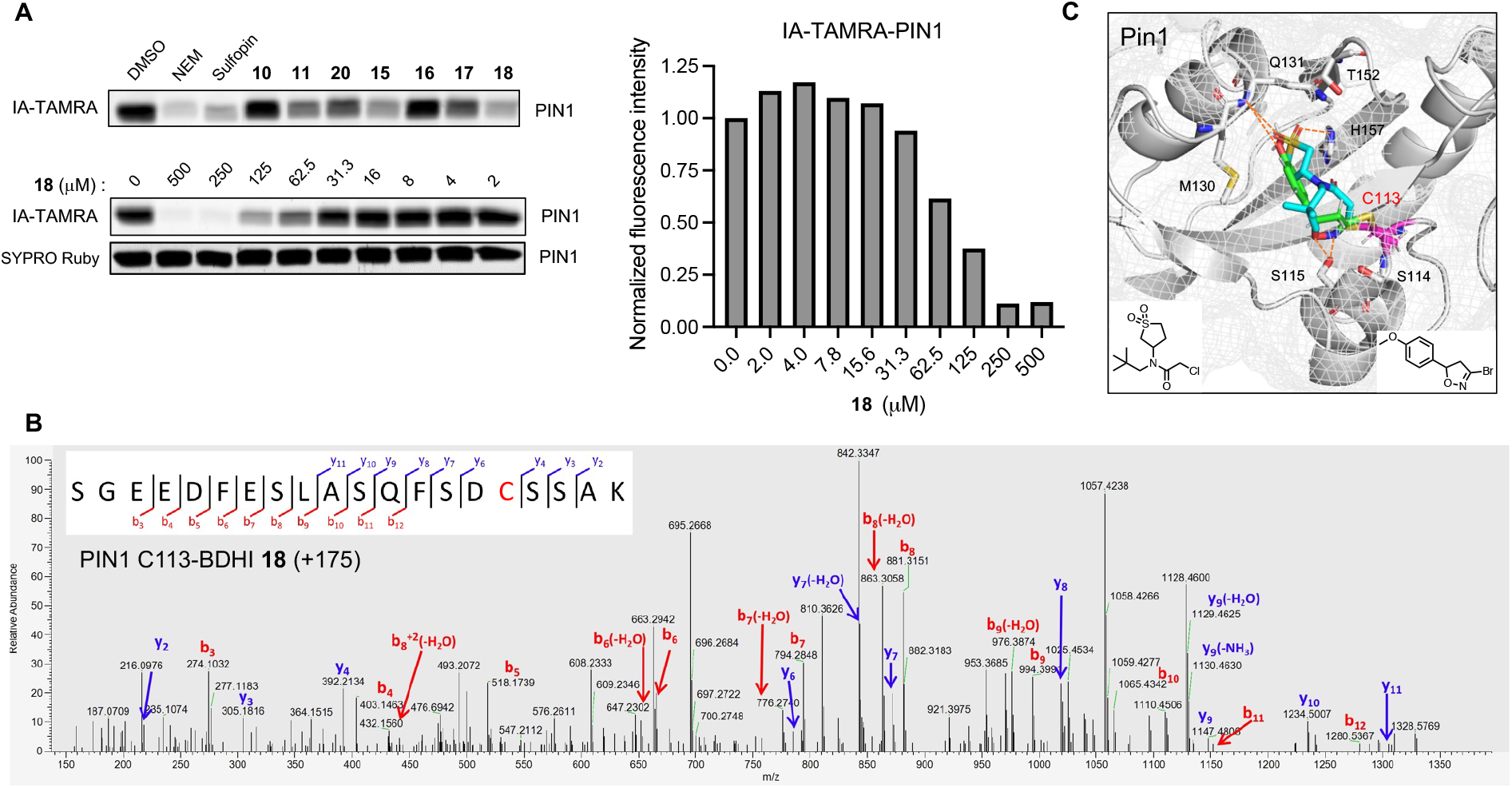
Validation of PIN1 as the BDHI-reactive protein. (A) Competitive and concentration-dependent inhibition of labeling of PIN1 by **18**. Recombinant PIN1 was preincubated with each compound (500 μM, 4h), labeled with IA-TAMRA (1 μM, 1 h) and subsequently analyzed by SDS-PAGE and in-gel fluorescence. (B) Annotated MS2 fragmentation spectra analysis of PIN1 liganded with BDHI **18** at the catalytic active site Cys113, highlighted in red. (C) Docking prediction of **18** binding to PIN 1 (PDB: 7EKV). The predicted binding pose of **18** (green) is superimposed with the crystal structure of PIN with a covalently bound inhibitor sulfopin (cyan) (PDB: 6VAJ).

## CONCLUSIONS

Covalent modification of proteins by small molecule electrophiles has been widely validated in drug discovery, protein engineering, imaging, and chemical proteomic applications. However, there remains a need for novel reactive functionalities that possess selectivity suitable for ligand discovery and pharmacological optimization. In this study, we used a computationally-guided approach to design and characterize a library of covalent fragments functionalized with a natural product-inspired 3-bromo-4,5-dihydroxazole (BDHI) electrophile. In vitro studies as well as proteome-wide competitive profiling experiments revealed that BDHIs engage a restricted subset of reactive cysteine residues in the human proteome. BDHI was found to possess tempered chemical reactivity relative to haloacetamide electrophiles, and our data implicate the identity of the reversible binding element, the composition and geometry of the complementary protein binding sites, and stabilization of the covalent intermediate as critical factors driving covalent cysteine labeling by this chemotype. Our studies also demonstrated the ability of BDHI-containing molecules to functionally regulate protein activity, for example via engagement of PIN1 and GSTP1. Finally, we highlight some limitations of our study and future work that is being performed to address them. First, the focus of our initial study was on the design and reactivity of this natural product-inspired electrophile rather than biological screening. However, in preliminary studies we have found that BDHI **18** can inhibit surface expression of activation markers (CD25 and CD69) and reduce cytokine secretion (IFN-γ, IL-2, and IL-6) from primary human T-cells while maintaining cell viability (Figure S10), suggesting the cell permeability of this electrophile. In the future, we anticipate the chemoproteomic probes developed here will enable in situ analysis of target engagement by **18** as well as other biologically active BDHI molecules. Second, the potencies of the molecules characterized in this study are relatively modest (micromolar range) compared to many covalent small molecules. Medicinal chemistry optimization of BDHI probes or, alternatively, substituting BDHI as warhead in known covalent drugs (e.g. ibrutinib) may provide a strategy to understand the pharmacological properties of this electrophile as well the interplay between non-covalent and covalent interactions. Related to this, acivicin has been reported to form a covalent bond with an active site threonine residue in *E. coli* γ-glutamyltranspeptidase (GGT).^12^ While our studies indicate BDHI-labeling of TG2 and ALDH1A1 is likely cysteine-directed, in the future it will be interesting to assess whether tighter-binding BDHIs may modify other amino acid targets. A unique property of the BDHI electrophile not explored in this study is the presence of a stereocenter. We anticipate the development of synthetic/purification routes to enantiomerically pure BDHIs may provide a facile strategy for differentiating new ligandable sites in proteins. Overall, our studies demonstrate how natural products, computation, and chemoproteomic profiling can augment electrophile design, providing a foundation for inhibitor discovery and optimization.

## ASSOCIATED CONTENT

### Supporting Information

The Supporting Information is available free of charge on the Publications website.

- Additional figures, tables, detailed procedures for all experiments, specifics for reagents and instruments used, and synthetic details (PDF)
- Table S2 proteomics data (XLSX)

## Notes

The authors declare no competing financial interest.

## ACKNOWLEDGMENT

The authors thank Dr. Jordan L. Meier (NCI) for the critical reading of the manuscript. This work was supported by the Intramural Research Program of the National Institutes of Health, National Cancer Institute, the Center for Cancer Research (ZIABC011961). This project was also funded in part under contract No. HHSN261200800001E. Part of this work was carried out on the MX1 and MX2 beamlines at the Australian Synchrotron, part of the Australian Nuclear Science and Technology Organization, and made use of the Australian Cancer Research Foundation detector on the MX2 beamline. We thank the beamline staff for their assistance. We acknowledge use of the Mass Spectrometry and Proteomics Facility, Bio21 Institute, The University of Melbourne. Funding from the Victorian Government Operational Infrastructure Support Scheme to St Vincent’s Institute is acknowledged. MWP is a National Health and Medical Research Council (NHMRC) of Australia Investigator (APP1117183).

## Notes

### Competing Interest Statement

The authors have declared no competing interest.

