## Supporting Information for "Covalent inhibition by a natural product-inspired latent electrophile"

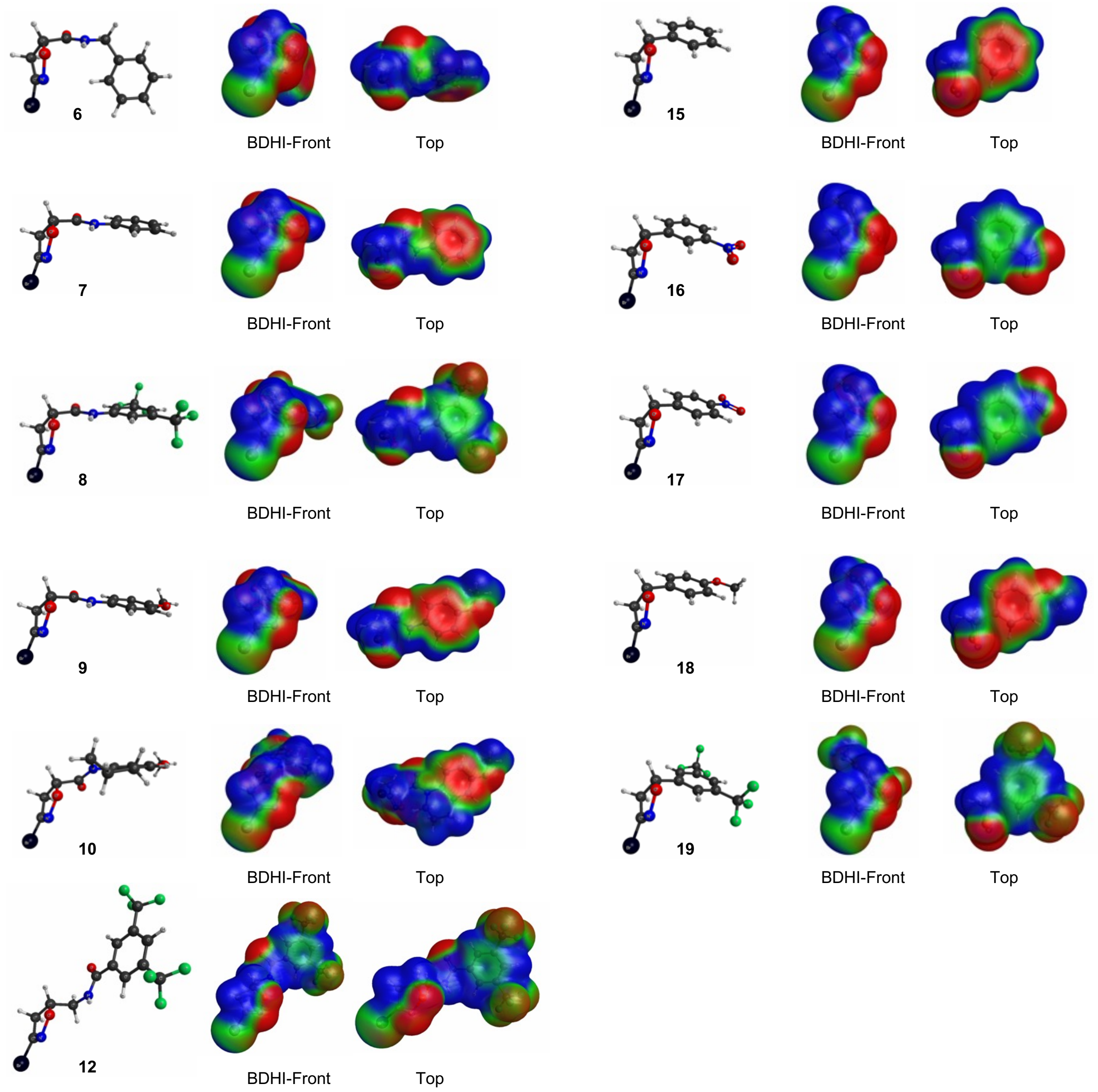


Figure S1. Molecular electrostatic potential surfaces of BDHI-containing fragments.

Table S1. Atomic charges calculated for BDHI ring

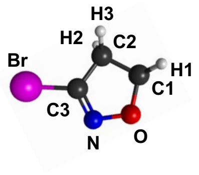


| | C1 | H1 | C2 | H2 | H3 | C3 | Br | N | O |
|---|---|---|---|---|---|---|---|---|---|
| **6** | 0.137 | 0.093 | -0.157 | 0.081 | 0.104 | 0.316 | -0.067 | -0.315 | -0.188 |
| **7** | 0.201 | 0.061 | -0.149 | 0.081 | 0.091 | 0.323 | -0.075 | -0.305 | -0.211 |
| **8** | 0.175 | 0.078 | -0.147 | 0.081 | 0.099 | 0.313 | -0.057 | -0.302 | -0.203 |
| **9** | 0.176 | 0.068 | -0.137 | 0.079 | 0.090 | 0.315 | -0.074 | -0.303 | -0.205 |
| **10** | 0.388 | 0.006 | -0.410 | 0.127 | 0.130 | 0.412 | -0.080 | -0.396 | -0.122 |
| **12** | 0.329 | 0.030 | -0.231 | 0.064 | 0.125 | 0.271 | -0.063 | -0.282 | -0.216 |
| **15** | 0.394 | 0.019 | -0.132 | 0.059 | 0.071 | 0.280 | -0.082 | -0.285 | -0.241 |
| **16** | 0.368 | 0.033 | -0.192 | 0.075 | 0.095 | 0.290 | -0.066 | -0.278 | -0.226 |
| **17** | 0.406 | 0.029 | -0.191 | 0.075 | 0.094 | 0.290 | -0.063 | -0.286 | -0.237 |
| **18** | 0.456 | 0.008 | -0.193 | 0.073 | 0.082 | 0.310 | -0.086 | -0.300 | -0.242 |
| **19** | 0.358 | 0.038 | -0.186 | 0.075 | 0.098 | 0.275 | -0.058 | -0.273 | -0.224 |

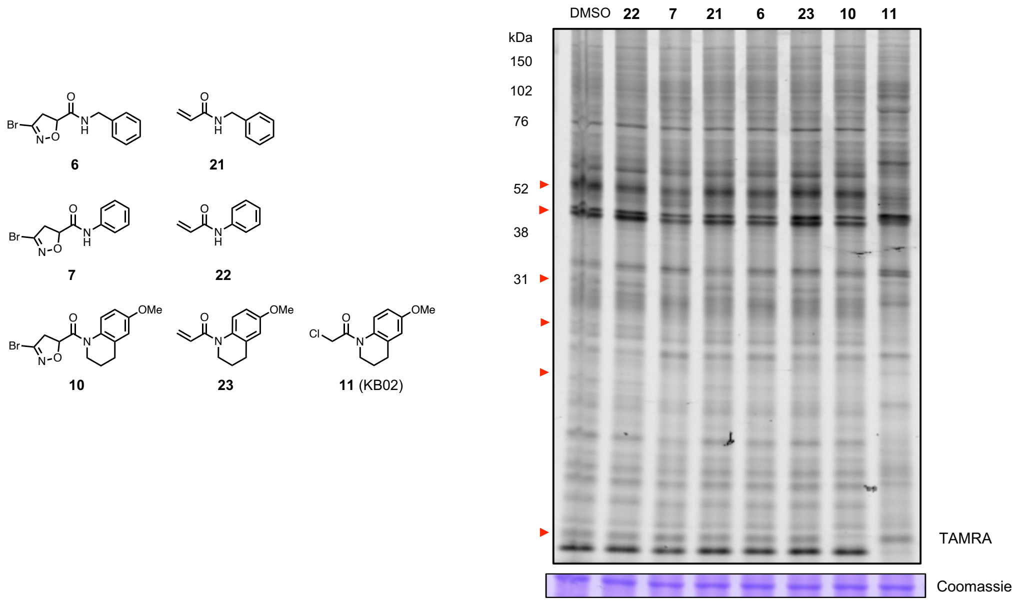


Figure S2. In-gel fluorescence image depicting competitive blockade of IA-TAMRA (1 μM, 1 h) labeling of proteins in Jurkat cell lysate by DMSO, KB02, acrylamide- or BDHI-containing fragments (500 μM, 4 h). Red arrows highlight protein bands that showed diminished IA-TAMRA labeling.

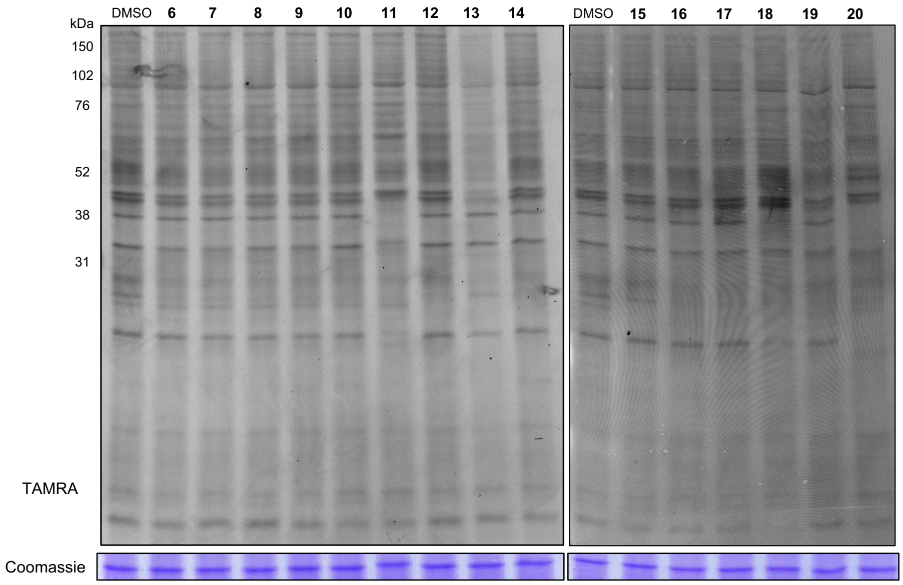


Figure S3. BDHI electrophile-containing fragments competitively block the labeling of proteins by BDHI probe. Soluble proteome from Jurkat cells was treated with the indicated fragments (500 μM, 4 h), followed by labelling with BDHI-TAMRA **2** (100 μM, 2 h) and analyzed by SDS-PAGE and in-gel fluorescence scanning.

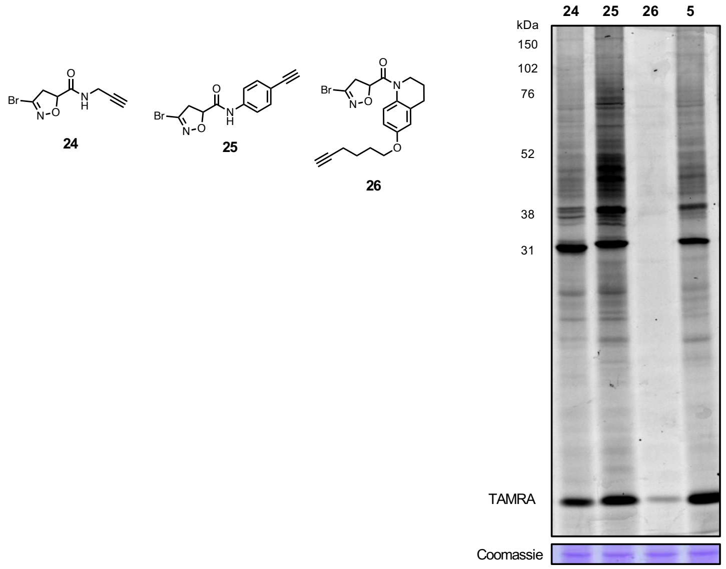


Figure S4. In-gel fluorescence image depicting protein bands labeled with BDHI-alkyne probes **24-26** in the human proteome. Soluble proteome from Jurkat cells was treated with the indicated probes at 500 μM for 2 h and subsequently subjected to CuAAC with a TAMRA-$N_3$, followed by SDS-PAGE and in-gel fluorescence scanning.

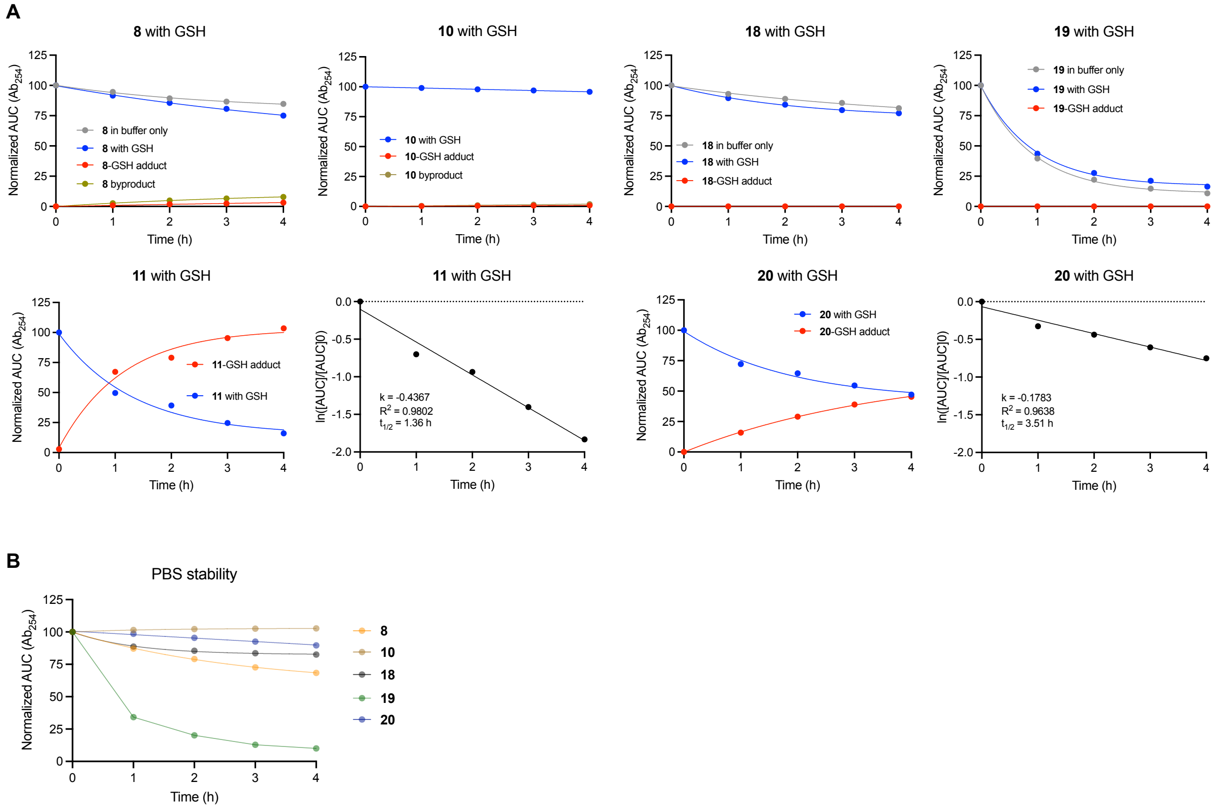


Figure S5. Solution reactivity and stability of BDHI analogs. (A) Relative compound consumption (blue) and product formation (red) measured in reactions with GSH at pH 7.4 by LC-MS. (B) Relative stability of compounds measured in PBS.

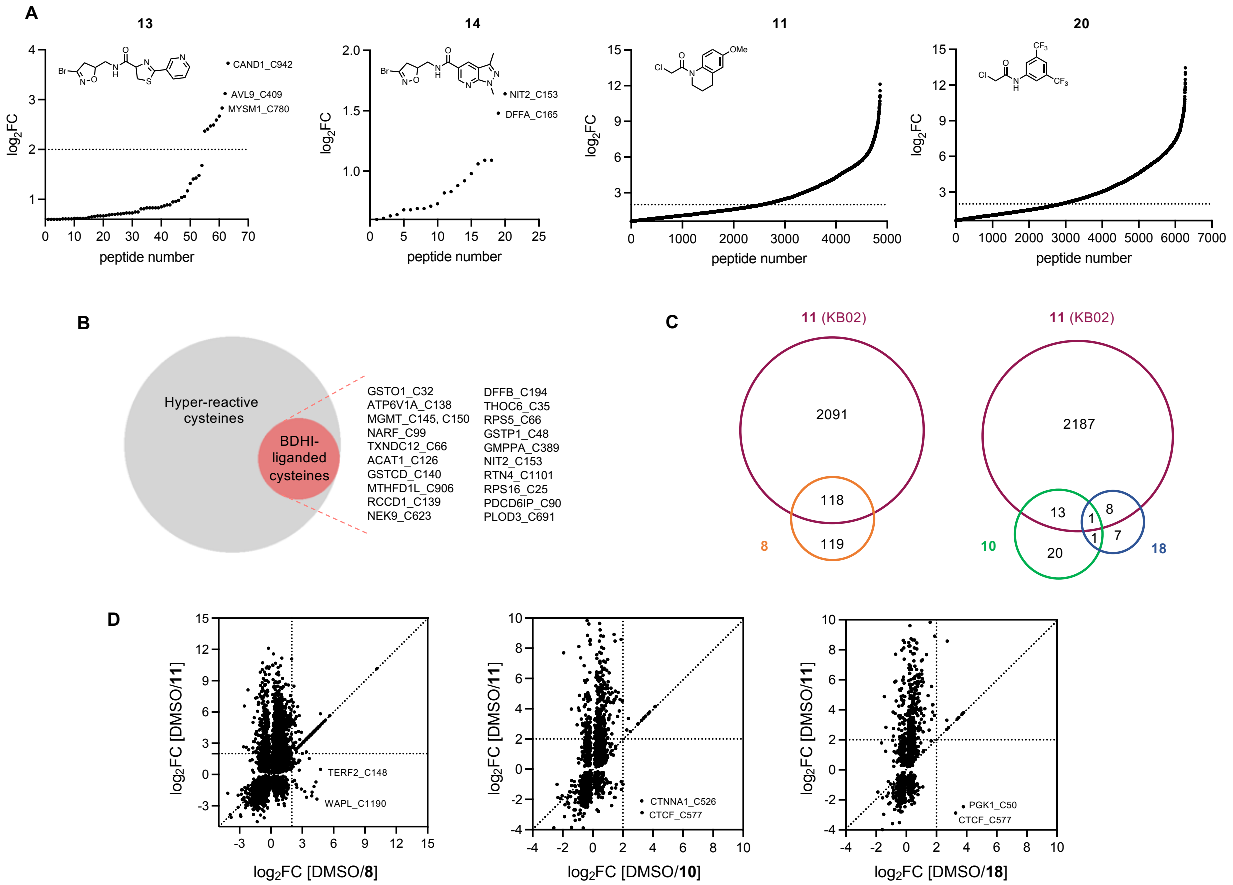


Figure S6. Competitive MS-based reactivity profiling defines cysteines liganded by BDHI analogs. (A) Structures and competition ratios ($log_2FC$ values) from Jurkat cell proteome treated with BDHI **13** and **14**, and CA **11** and **20**. (B) Venn diagram representation of number of hyper-reactive cysteines and BDHI-liganded cysteines. (C) Venn diagram representation of number of cysteine sites significantly liganded ($log_2FC > 2$) by BDHIs in comparison to KB02 (**11**). Results were obtained by comparing the site overlap at a given competition ratio threshold for each ligand. (D) Scatter plot of competition ratios for quantified cysteine-containing peptides ($p$ value < 0.05) targeted by KB02 (**11**) and each BDHI ligand **8**, **10**, and **18**.

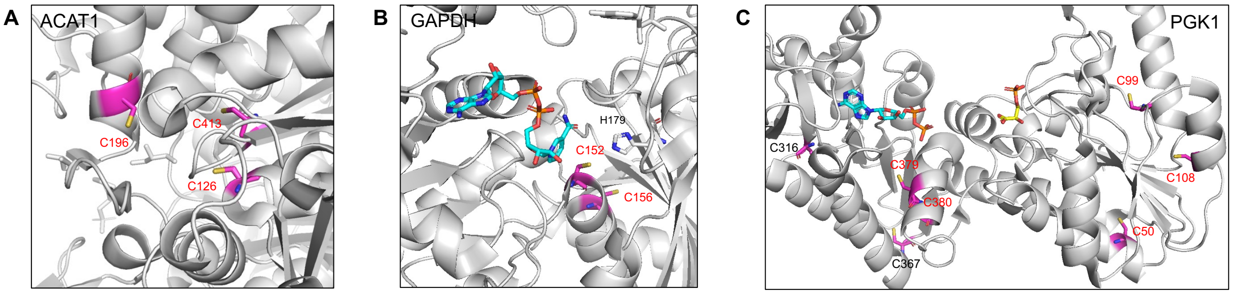


Figure S7. (A) Crystal structure of ACAT1 highlighting the active-site cysteine residues (PDB: 2IB8). (B) Crystal structure of GAPDH with $NAD^+$ bound (PDB: 1U8F). (C) Crystal structure of PGK1 depicting reactive cysteines (PDB: 2XE7).

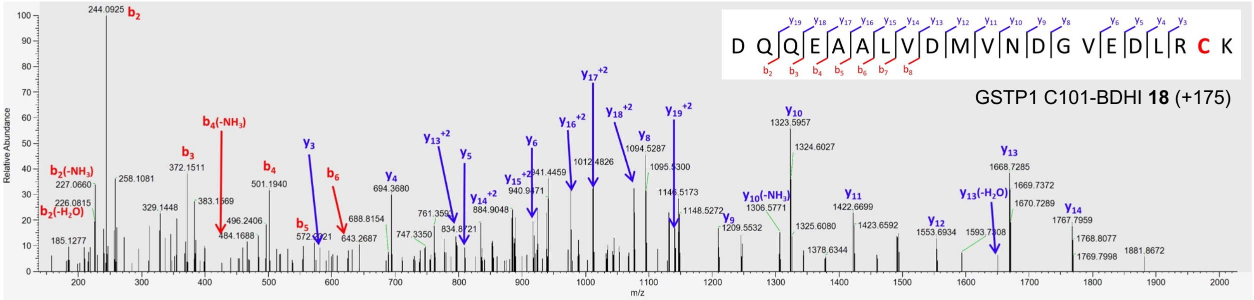


Figure S8. Binding site analysis of BDHI **18** with GSTP1 at Cys101.

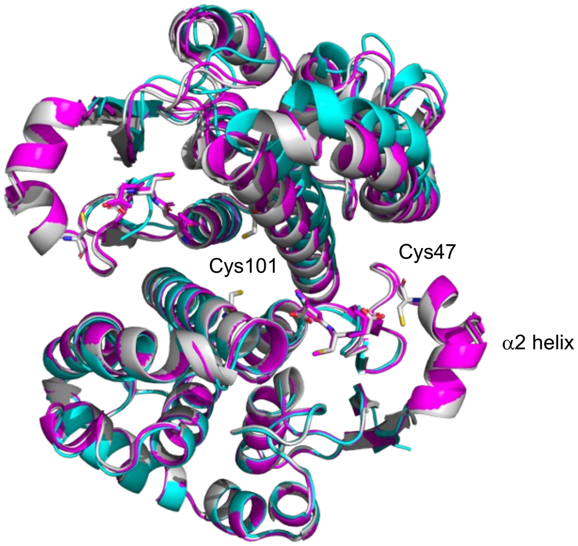


Figure S9. Structures of GSTP1. Cyan: after soaking with 5 mM **18** for 2 h at pH 7; Pink: after soaking with 5 mM **18** for 2 h at pH 7 and back-soaking with 10 mM GSH; Grey: crystallized with GSH (PDB: 5GSS). The GSH molecules and residues Cys47 and Cys101 are in stick representation.

Table S3. Summary of observations from structures of soaked crystals

| Fragment | Soaking Concentration | Time | Soak 2 | Crystal ID | Electron density in $\alpha 2$ helix (monomer A) | Electron density in $\alpha 2$ helix (monomer B) |
|---|---|---|---|---|---|---|
| DMSO | 5% | 2 h | – | ASP0095_07_0200<br>ASP0095_08_0202 | no loss | no loss |
| **18** | 1 mM | 2 h | – | ASP0095_04_0194<br>ASP0095_05_0196 | no loss | no loss |
| **18** | 5 mM | 2 h | – | ASP0220_03_0102<br>ASP0095_01_0188*<br>ASP0095_02_0190*<br>ASP0095_03_0192* | complete loss<br>significant loss<br>no loss<br>complete loss | complete loss<br>significant loss<br>minor loss<br>complete loss |
| **18** | 0.5 mM | 74h | – | ASP0220_15_0219<br>ASP0220_16_0221 | minor loss | significant loss |
| **18** | 2.5 mM | 3 weeks | – | ASP0223_05_0063 | complete loss | complete loss |
| DMSO | 5% | 2 h | 10 mM GSH 1 h | ASP0220_11_0204 | no loss; GSH bound | no loss; GSH bound |
| **18** | 1 mM | 2 h | 10 mM GSH 1 h | ASP0220_12_0206 | no loss; GSH bound | no loss; GSH bound |
| **18** | 5 mM | 2 h | 10 mM GSH 1 h | ASP0220_13_0208* | no loss; GSH bound | no loss; GSH bound |
| DMSO | 5% | 2 h | 10 mM GSH 2 h | ASP0223_01_0209<br>ASP0223_02_0214 | no loss; GSH bound | no loss; GSH bound |
| **18** | 1 mM | 2 h | 10 mM GSH 2 h | ASP0223_03_0216<br>ASP0223_04_0218<br>ASP0223_05_0220 | no loss; GSH bound | no loss; GSH bound |
| **18** | 5 mM | 2 h | 10 mM GSH 2 h | ASP0223_06_0222*<br>ASP0223_08_0226 | no loss; GSH bound | no loss; GSH bound |

*crystals also subjected to intact mass spectrometric analysis

Table S4. Relative abundance of GSTP1 species observed in GSTP1 crystals soaked with **18** alone or with GSH back-soak

| Crystal | % apoGSTP1 | % GSTP1-fragment | % GSTP1-(fragment)$_2$ |
|---|---|---|---|
| ASP0095_01_0188 | 30 | 29 | 30 |
| ASP0095_02_0190 | 29 | 28 | 32 |
| ASP0095_03_0192 | 28 | 31 | 29 |
| ASP0220_13_0208 | 28 | 27 | 33 |
| ASP0223_06_0222 | 21 | 29 | 37 |

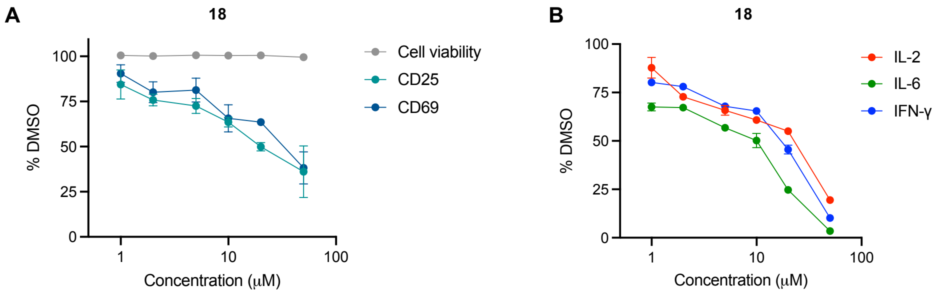


Figure S10. BDHI analog **18** suppresses T cell activation. Concentration dependent effect of BDHI **18** on T cell activation (*n* = 4) measured by surface expression of CD25 and CD69 (A) and secretion of cytokines IL-2, IL-6, and IFN-γ (B). Primary human T cells were treated with **18** under TCR-stimulating conditions (96-well plates pre-coated with αCD3 and αCD28) for 24 h.

#### Materials and Methods

##### Cell culture and isolation of proteome

All cell lines were cultured at 37°C under 5% $CO_2$ atmosphere. Jurkat cells were cultured in RPMI supplemented with 10% fetal bovine serum (FBS), 1% penicillin/streptomycin (PS), and 2 mM L-glutamine, while HeLa and HEK293T were cultured in DMEM supplemented likewise as described above. Whole blood samples from healthy adult donors were obtained from the occupational health service, National Cancer Institute (NCI), Frederick. Peripheral blood mononuclear cells (PBMCs) were isolated by Ficoll-paque (Sigma-Aldrich) density gradient centrifugation (400 *g*, 30 min, room temperature). The cells were pelleted (200 *g*, 5 min) and washed twice with PBS. Total T cells were isolated by negative selection using human T cell isolation kit (Invitrogen, 11344D) according to manufacturer's instructions. For preparation of proteomes, cells were harvested, pelleted (3000 *g*, 5 min, 4°C), and washed with ice-cold PBS. The cell pellets were resuspended in ice-cold PBS (37.5 μL per 1 x $10^6$ cells), lysed by sonication using a 700-W QSonica Q700 sonicator (7 x 3 sec pulse, amplitude 1, 30 sec resting on ice between pulses), and pelleted by ultracentrifugation (15000 *g*, 20 min, 4°C). The supernatant was collected, filtered through a 0.45 μm syringe filter, and quantified using a Pierce BCA protein assay (Thermo Scientific, 23227). Protein concentrations were adjusted to 1.5 - 2 mg/mL and stored at -80°C.

##### Solution reactivity assay with relevant nucleophiles by LC-MS

To 98 μL of 5 mM nucleophile [GSH: 1/10 dilution of freshly prepared 50 mM GSH in assay buffer (9:1 mixture of 0.1 M pH 7.4 phosphate buffer and acetonitrile); Boc-Cys-OMe: 1/100 dilution of 500 mM DMSO stock in assay buffer] was added 1 μL of compound (10 mM in DMSO, final concentration: 500 μM) and 1 μL of internal standard (10 mM rhodamine B or rhodamine 110 in DMSO, final concentration: 500 μM) (internal standard chosen such that its $UV_{254}$ does not overlap with tested compound). The resulting reaction mixture was analyzed by LC/MS at 0, 1, 2, 3, 4, and 16 h time points at room temperature. LC/MS runs were performed on Agilent 6250 Accurate – Mass Q-TOF LC/MS with Agilent Poroshell 5 mm 300SB-C18 column. 0.1% formic acid in water and 0.1% formic acid in acetonitrile were used as mobile phases A and B, respectively, with flow rate of 1.0 mL/min. The gradient used was 5% to 100% B from 0 to 4 min and holding at 100% B for 1 min. The analysis was done on Agilent MassHunter Qualitative Analysis B.07.00. Abundance was determined by calculating the area under the curve (AUC) of absorbance at 254 nm and normalizing to the internal standard.

##### Competitive gel-based protein profiling

**Using IA-TAMRA:** Jurkat cell lysates (20 μL, 1 mg/mL in PBS) were treated with DMSO or compound (1 μL of 10 mM in DMSO, final concentration: 500 μM) at 37°C for 4 h, followed by treatment with IA-TAMRA (1 μL of 20 μM in DMSO, final concentration: 1 μM) at 37°C for 1 h. Reactions were quenched by adding 4x reducing Laemmli SDS sample buffer (6.5 μL), then samples were boiled at 100°C for 5 min, cooled to room temperature, and resolved via SDS-PAGE on a 12% polyacrylamide gel. In-gel fluorescence scanning was performed using a Typhoon FLA 9500 (GE Healthcare).

**Using click chemistry with TAMRA-$N_3$:** Jurkat cell lysates (50 μL, 1 mg/mL in PBS) were treated with DMSO or compound (0.5 μL of 100x in DMSO) at 37°C for 4 h, followed by treatment with desired alkyne probe (0.5 μL of 100x in DMSO) at 37°C for 1 h. Labeled proteins were subjected to copper-catalyzed azide-alkyne cycloaddition (CuAAC) reaction by adding 6 μL of click mixture consisting of the following: 3 μL of 1.7 mM TBTA ligand in 4:1 *t*-BuOH/DMSO, 1 μL of 50 mM $CuSO_4$ in water, 1 μL of 50 mM TCEP-HCl in water (freshly prepared), and 1 μL of 12.5 mM TAMRA-azide in DMSO. The samples were incubated at room temperature for 1 h. Reactions were quenched by adding 4x reducing Laemmli SDS sample buffer (16.5 μL), then samples were boiled at 100°C for 5 min, cooled to room temperature, and resolved via SDS-PAGE on a 12% polyacrylamide gel. In-gel fluorescence scanning was performed using a Typhoon FLA 9500 (GE Healthcare).

##### Competitive chemoproteomic profiling with IA-DTB

**Compound pretreatment and IA-DTB labeling.** DMSO control and compound-treated samples were prepared in triplicate. Jurkat cell lysates (100 μL, 1 mg/mL in PBS) were incubated with DMSO or compound (1 μL of 50 mM in DMSO, final concentration: 500 μM) for 4 h at 37°C, followed by treatment with desthiobiotin-iodoacetamide (IA-DTB, 1 μL of 50 mM in DMSO, final concentration: 500 μM) at 25°C in the dark with shaking for 1 h. 2.5 μL of freshly prepared DTT (31 mg/mL in water) were added and incubated at 25°C in the dark for 30 min. Free thiols were alkylated by incubating samples with 5 μL of 2-iodoacetamide (74 mg/mL in water) at 25°C in the dark for 30 min. Proteins were precipitated by adding 500 μL of 4:1 MeOH/$CHCl_3$ and 300 μL of water, thoroughly vortexing, then centrifuging (18,000 *g*, 2 min, 25°C) and removing the top aqueous layer without disturbing the protein disc at the layer interface. 500 μL of MeOH were added and vortexed thoroughly, centrifuged (18,000 *g*, 2min, 25°C), and the supernatant was removed (2x). Samples were quickly dried using a SpeedVac, and the protein pellets were stored at -20°C until further use.

**Digestion and streptavidin enrichment.** Each sample pellet was resuspended in 200 μL of EasyPep lysis buffer (Thermo, A45735) supplemented with Trypsin/LysC (Thermo, A40007) at a concentration of 27 ng/μL. Samples were incubated at 37°C with shaking for 24 h after which point 50 μL of 10 μg/mL TMTpro 18-plex (Thermo, A52045) reagent was added to each sample and incubated for 1 h at 25°C with shaking. Excess TMTpro was quenched with 50 μL of 5% hydroxylamine, 20% formic acid for 10 min and samples were then combined. Samples were cleaned using EasyPep Maxi columns (Thermo, A45734) as described in the manual. Eluted peptides were dried in speed-vac. High-Capacity Streptavidin Agarose (HCSA, Thermo, 20359) was used for enrichment of DTB peptides. In a clean 1.5 mL tube, 400 μL of HCSA slurry was added (50% slurry, 200 mg of settled beads) and centrifuged and the liquid was removed and the beads were washed three times with 400 μL of PBS. The dried peptides were resuspended in 1 mL PBS and added to the beads and mixed with end-over-end rotation for 1 h at room temperature. The slurry was transferred to a micro spin column (Thermo, 69705) and centrifuged to remove supernatant. The beads were then washed with 500 μL of 0.1% NP-40 in PBS, PBS and water (five times each). Bound peptides were eluted with 300 μL of 50% ACN, 0.5% TFA (3x) and then all three elution fractions were combined and dried.

**LC/MS analysis of enriched peptides.** All samples were analyzed on a Dionex U3000 RSLC in front of a Orbitrap Eclipse (Thermo) equipped with an EasySpray ion source with FAIMS™ where indicated. Advanced Peak Determination, Monoisotopic Precursor selection (MIPS), and EASY-IC for internal calibration were enabled and dynamic exclusion was set to a count of 1 for 15 sec in all methods. Solvent A consisted of 0.1% formic acid (FA) in water and Solvent B consisted of 0.1% FA in 80% acetonitrile. Loading pump consisted of Solvent A and was operated at 7 mL/min for the first 6 min of the run then dropped to 2 mL/min when the valve was switched to bring the trap column (Acclaim™ PepMap™ 100 C18 HPLC Column, 3 mm, 75 mm I.D., 2 cm, PN 164535) in-line with the analytical column EasySpray C18 HPLC Column, 2 mm, 75 mm I.D., 25 cm, PN ES902). Enriched peptides were resuspended in 40 mL of 0.1% FA and 12 μL was analyzed in triplicate. Each of the three injections consisted of the same LC gradient conditions and global MS parameters, with only the FAIMS compensation voltages (CVs) changing for each method. Each run used a linear LC gradient of 5-7% B for 1 min, 7-30% B for 34 min, 30-50% B for 15 min, 50-95% B for 4 min, holding at 95% B for 7 min, then re-equilibration of analytical column at 300 nL/min at 5% B for 17 min. All three MS injections employed the TopSpeed method with three FAIMS compensation voltages (CVs) and a 1 second cycle time for each CV (3 second cycle time total) that consisted of the following: Spray voltage was 2200V and ion transfer temperature was 300°C. MS1 scans were acquired in the Orbitrap with resolution of 120,000, AGC of 4e5 ions, and max injection time of 50 ms, mass range of 350-1600 m/z; MS2 scans were acquired in the Orbitrap using TurboTMT method with resolution of 15,000, AGC of 1.25e5, max injection time of 22 ms, HCD energy of 30%, isolation width of 1.6 Da, intensity threshold of 2.5e4 and charges 2-5 for MS2 selection. Advanced Peak Determination, Monoisotopic Precursor selection (MIPS), and EASY-IC for internal calibration were enabled and dynamic exclusion was set to a count of 1 for 15 sec in all methods. The only difference in the methods was the CVs used, Method 1 used CVs of -45, -60, -75; method 2 used CVs of -50, -65, -80; method 3 used CVs of -55, -70, -85.

**Database search and post-processing analysis.** All data were searched in Proteome Discoverer 2.4 using the Sequest node. Data was searched against the Uniprot Human database from Feb 2020 using a full tryptic digest, 2 max missed cleavages, minimum peptide length of 6 amino acids and maximum peptide length of 60 amino acids, an MS1 mass tolerance of 10 ppm, MS2 mass tolerance of 0.02 Da, fixed modifications for TMTpro (+304.207) on lysine and peptide *N*-terminus, variable oxidation on methionine (+15.995 Da), and variable carbamidomethyl (+57.021) and desthiobiotin iodoacetamide (+296.185) on cysteine. Percolator was used for FDR analysis and IMP-ptmRS for site localization. TMTpro reporter ions were quantified using the Reporter Ion Quantifier node and normalized on total peptide amount. For the final group comparisons only peptides that were observed in 6 or more total samples were included.

**IA-TAMRA labeling of recombinant proteins**

Recombinant protein (diluted in 20 μL of PBS) was treated with DMSO or compound (1 μL of 10 mM in DMSO, final concentration: 500 μM) and incubated at 37°C for 4 h, followed by treatment with IA-TAMRA (1 μL of 20 μM in DMSO, final concentration: 1 μM) for 1 h. Reactions were quenched by adding 4x reducing Laemmli SDS sample buffer (6.5 μL), then samples were boiled at 100°C for 5 min, cooled to room temperature, and resolved via SDS-PAGE on a 12% polyacrylamide gel. In-gel fluorescence scanning was performed using a Typhoon FLA 9500 (GE Healthcare).

**Probe 5 labeling of recombinant proteins**

Recombinant protein (diluted in 20 μL of PBS) was treated with DMSO or compound (1 μL of 10 mM in DMSO, final concentration: 500 μM) and incubated at 37°C for 4 h, followed by treatment with **5** (1 μL of 10 mM in DMSO, final concentration: 500 μM) for 2 h. Labeled proteins were subjected to copper-catalyzed azide-alkyne cycloaddition (CuAAC) reaction by adding 2.4 μL of click mixture consisting of the following: 1.2 μL of 1.7 mM TBTA ligand in 4:1 *t*-BuOH/DMSO, 0.4 μL of 50 mM $CuSO_4$ in water, 0.4 μL of 50 mM TCEP-HCl in water, and 0.4 μL of 12.5 mM TAMRA-azide in DMSO. The samples were incubated at room temperature for 1 h. Reactions were quenched by adding 4x reducing Laemmli SDS sample buffer (6.5 μL), then samples were boiled at 100°C for 5 min, cooled to room temperature, and resolved via SDS-PAGE on a 12% polyacrylamide gel. In-gel fluorescence scanning was performed using a Typhoon FLA 9500 (GE Healthcare).

**GSTP1 enzymatic assay**

rhGSTP1 was adjusted to 2.2 ng/μL in assay buffer (0.1 M phosphate buffer, pH 7.0). In separate tubes, samples were prepared in duplicate by combining 19 μL of adjusted rhGSTP1 (2.2 ng/μL) and 1 μL of either DMSO or 20X compound in DMSO. A blank sample was also included by combining 19 μL of assay buffer and 1 μL of DMSO. Samples were incubated at room temperature for 15 min then transferred to a 96 well plate. 20 μL of 4 mM GSH in assay buffer (prepared from 250 mM stock in $ddH_2O$) were added, followed by 40 μL of 2 mM 1-bromo-2,4-dinitrobenzene (BDNB) substrate in assay buffer (prepared from 75 mM stock in ethanol). Absorbance was measured at 340 nm for 10 min in 30 sec intervals using a BioTek Cytation 5 plate reader.

**GSTP1 crystallography**

Full-length human GSTP1 was purified from *E. coli* by affinity chromatography. The protein was stored in 6 mg/mL aliquots in 10 mM potassium phosphate pH 6.5 and 0.1 mM EDTA at -80°C.

**Crystallization.** Protein was thawed and diluted to 5 mg/mL in the absence of ligands (apoGSTP1). Crystallization drops were prepared in hanging drop plates, with 1 μL of protein solution added to 1 μL of crystallization buffer containing 0.1 M MES pH 6.0, 20 mM $CaCl_2$, 26% PEG8000, at room temperature. After 1 h, drops were streak-seeded using a cat whisker dipped in drops containing apoGSTP1 crystals previously grown in crystallization buffer plus 10 mM DTT.

**Crystal soaks.** Soaking solutions were prepared containing crystallization buffer with 1 mM or 5 mM **5** or **18**. An equal volume of soaking solution was added to crystallization drops containing apoGSTP1 crystals with crystals soaked for 74 h or three weeks. Alternatively, apoGSTP1 crystals were transferred into buffer containing either 0.1 M MES pH 6 or 7, 20 mM $CaCl_2$, 26% PEG8000 and 1 mM or 5 mM fragment. Back-soaking with GSH was also attempted. After soaking with 1 mM or 5 mM **18**, an equal volume of soaking solution containing GSH (0.1 M MES pH 6, 20 mM $CaCl_2$, 26% PEG8000 and 20 mM GSH) was added to the drop.

**Crystal harvesting.** Crystals were transferred into cryobuffers containing 0.1 M MES pH 6, 20 mM $CaCl_2$, 26% PEG8000 ± 500 μM **5** or **18** and 5% hexylene glycol, then 10% hexylene glycol. Crystals were flash frozen and stored in liquid nitrogen.

**Crystallography.** Data were collected at 100 K at the Australian Synchrotron's MX1[1]or MX2[2] beamline, with crystals diffracting to resolutions of 1.99-2.71 Å in the *C2* space group. Data were integrated using XDS[3] and scaled with AIMLESS from the CCP4i2 program suite.[4] DIMPLE (http://ccp4.github.io/dimple/) was used to generate maps and a rigid body refinement was performed in REFMAC5[5] using PDB 5DCG[6] (with heteroatoms removed) or 5GSS[7] (with GSH and waters) as search models. An additional round of model building and refinement was performed using Coot[8] and REFMAC5, as needed.

**GSTP1 crystals for mass spectrometry.** GSTP1 crystals were harvested from crystal loops after X-ray diffraction experiments. The crystals had been soaked for 2 h at pH 7 in 5 mM **18** only or subsequently back-soaked with 10 mM GSH. Crystals were transferred from the soaking solution into two cryobuffer solutions where no fragment was present and frozen in liquid nitrogen. After diffraction, crystals were thawed and transferred into acetonitrile, then MilliQ water. After failing to dissolve in water, 6 M urea was added to dissolve the crystals. The 2-4 μL drop was then diluted with 10 μL of MilliQ water for analysis. Dissolved protein crystals were analysed for total protein mass by electrospray ionisation mass spectrometry on a Acquity UHPLC H-Class/Waters Vion IMS-QTOF system (Melbourne Mass Spectrometry and Proteomics Facility, Bio21 Institute, University of Melbourne). Data were analysed using MassLynx software.

**Modification site analysis of BDHI 18**
1 μg of recombinant protein was diluted in 40 μL of PBS, then BDHI **18** (1 μL of 10 mM in DMSO) were added and incubated at 37°C for 4 h in triplicate, then samples were flash frozen and stored at -80°C until further use.

**Digestion of GSTP1 and PIN1.** Each replicate was treated with 100 μL of EasyPep Lysis buffer, 50 μL of reducing solution and 50 μL of alkylating solution provided with the EasyPep kit (Thermo, A45733). Samples were incubated at 25°C for 1 h in the dark then treated with 10 μL of 100 ng/μL of trypsin/LysC (Thermo, A40009) and incubated at 37°C overnight ~18 h. Each sample was treated with 50 μL of stop solution provided with the EasyPep kit and cleaned using EasyPep 96-well plate, eluted in 300 μL and dried in speedvac.

**LC/MS analysis.** All samples were analyzed on a Dionex U3000 RSLC in front of a Orbitrap Eclipse (Thermo) equipped with an EasySpray ion source with FAIMS™ interface. Advanced Peak Determination, Monoisotopic Precursor selection (MIPS), and EASY-IC for internal calibration were enabled and dynamic exclusion was set to a count of 1 for 15 sec in all methods. Solvent A consisted of 0.1% FA in water and Solvent B consisted of 0.1% FA in 80% ACN. Loading pump consisted of Solvent A and was operated at 7 μL/min for the first 6 minutes of the run then dropped to 2 μL/min when the valve was switched to bring the trap column (Acclaim™ PepMap™ 100 C18 HPLC Column, 3 μm, 75 μm I.D., 2 cm, PN 164535) in-line with the analytical column EasySpray C18 HPLC Column, 2 μm, 75 μm I.D., 25 cm, PN ES902). Digested peptides were resuspended in 50 μL of 0.1% FA and 5 μL was analyzed in triplicate at a flow rate of 300 nL/min using the same linear LC gradient of 5-7% B for 1 min, 7-30% B for 34 min, 30-50% B for 15 min, 50-95% B for 4 min, holding at 95% B for 7 min, then re-equilibration of analytical column at 5% B for 17 min. For each sample all three MS injections employed the TopSpeed method with three FAIMS compensation voltages (CVs) and a 1 second cycle time for each CV (3 second cycle time total) that consisted of the following: Spray voltage of 2200V and ion transfer temperature of 300°C. MS1 scans were acquired in the Orbitrap with resolution of 120,000, AGC of 4e5 ions, and max injection time of 50ms, mass range of 350-1600 m/z; MS2 scans were acquired in the Orbitrap using resolution of 15,000, AGC of 5e4, max injection time of 22 ms, HCD energy of 30%, isolation width of 0.4 Da, intensity threshold of 2.5e4 and charges 2-5 for MS2 selection. The only difference in the methods was the FAIMS CVs used.

**Database search general parameters.** All data were searched in Proteome Discoverer 2.4 using the Sequest node. For each sample the three MS files acquired were searched together to generate one output for each individual sample. Data was searched against the Uniprot Human database from Feb 2020 using a full tryptic digest, 2 max missed cleavages, peptide length of 6-40 amino acids, an MS1 mass tolerance of 10 ppm, MS2 mass tolerance of 0.02 Da, oxidation on methionine (+15.995 Da) and carbamidomethyl on cysteine (+57.021) were set to variable modifications on all searches. Additional variable modifications on cysteine of BDHI **18** (+175.063). FDR was calculated using Fixed Value PSM and modification sites were localized with the IMP-ptmRS node.

**Cell Titer Blue cell viability assay**
Cells were cultured as described above, and all incubation steps were performed at 37°C under 5% $CO_2$ atmosphere. 100 μL of cells were seeded into 96-well plates (Jurkat: 50,000 cells/well; HeLa/HEK293T: 10,000 cells/well; HeLa and HEK293T cells were left to adhere overnight while Jurkat cells were immediately treated with compounds). A blank was included using 100 μL of media containing no cells. Cells were treated with either DMSO or compounds (1 μL of 100X compound in DMSO). Following 24 h incubation, 20 μL of CellTiterBlue (Promega, G8081) were added and incubated for 2 h. Viability was

quantified by fluorescence (ex. 560 nm, em. 590 nm) using a BioTek Cytation 5 plate reader. Fluorescence values were normalized to DMSO (100% viability) and blank wells (0% viability). Experiments were performed in biological duplicate.

**T cell activation screen**
Tissue culture treated 96-well plates were coated with αCD3 (5 μg/mL) and αCD28 (2 μg/mL) antibodies from BioXcell in DPBS (200 μL/well) and left at 4°C overnight. Next day, the pre-coated plates were washed with complete media (3 x 200 μL). Freshly isolated total T cells were re-suspended in complete IMDM media supplemented with 10% FBS, penicillin (100 U/mL), and streptomycin (100 μg/mL) at $2 \times 10^6$ cells/mL. T cells were added to each well (200 μL/well, $4 \times 10^5$ cells/well), followed by the addition of 0.8 μL of 250x compound in DMSO to the respective wells. Cells were then incubated with the compound at 37°C in a 5% $CO_2$ containing incubator for 24 h. Following the treatment, the cells were collected for subsequent analysis. The supernatants were kept and stored at -20°C for the analysis of secreted cytokines.

**Flow cytometry analysis**
Following the PBS washes, the cells were stained with fixable near-IR LIVE/DEAD cell stain (Invitrogen, L34975) according to manufacturer's instructions. Briefly, LIVE/DEAD dye was diluted with PBS (1:1000) and added to each tube (200 μL) and the cells were incubated for 30 min at room temperature in the dark. The cells were then pelleted (500 *g*, 5 min), and washed with PBS (2 x 200 μL). The cells were incubated with mixture of antibodies for the appropriate cell surface markers (αCD25-Alexa Fluor 700 and αCD69-Alexa Fluor 647 antibodies, Biolegend) diluted in PBS containing 2% FBS. After 1 h incubation with antibodies on ice, cells were washed once with PBS, resuspended in PBS (200 μL), and analyzed by flow cytometry (BD Symphony).

**Bio-Plex quantification of secreted cytokines**
The supernatants collected from T cell activation assays in the presence or absence of compounds were used for the analysis of secreted cytokines using Bio-Plex Pro Human Cytokine assay (Bio-Rad) according to manufacturer's instructions. 50 μL of supernatant was used to detect different cytokines secreted during T cell activation and analyzed by Bioplex 200 system.

***In silico* modeling**
All *in silico* studies were conducted on ICM-Pro version 3.9 (Molsoft LLC, San Diego, CA). PDB structures were converted to internal coordinates by software's built-in function with water removal, hydrogen optimization and HisProAnsGlnCys optimization. Binding sites were determined by ICM pocket finder function, and target residue for covalent docking was selected based on either the proteomics LC/MS-MS data obtained in this study or based on reported literature. Reaction type was restricted to SNAR-type, where the bromine of the dihydroisoxazole moiety is replaced to a cysteine. PIN1 docking was performed using the PDB code 7EKV with a thoroughness of 500 and N conformation of 500. All docking was conducted in triplicate, unless stated otherwise. Best scoring pose was selected for visual inspection and analysis.

**Computational Chemistry**
Quantum chemistry calculations were performed the GAMESS package[9-10] where 6-31G(d) basis sets (spherical harmonics) were used.[11-13] Density functional theory (DFT)[14] utilizing the M11[15] functional was used to optimize geometries and subsequent atomic charges were computed by fitting to the electrostatic potentials via a geodesic point selection scheme.[16] Molecular electrostatic potential (MEP) surfaces were illustrated using MacMolPlt.[17]

**General chemical materials and methods**
All solvents were purchased from Fisher Scientific and Sigma Aldrich (HPLC grade). All reagents were purchased from Sigma Aldrich and Oakwood Chemicals. All water-sensitive reactions were performed in anhydrous solvents under positive pressure of nitrogen. Reactions were analyzed by LC-MS (Agilent). Reverse-phase HPLC was conducted with an Agilent 1260 infinity II using C18 columns (5 μm, 100 Å, 250 x 10 mm) (solvent A: water with 0.1% (v/v) TFA, solvent B: $CH_3CN$ with 0.1% (v/v) TFA). NMR spectra were recorded on a Varian 400 MHz (400/100) or Varian 500 MHz (500/125) equipped with a pulsed field gradient

accessory. Chemical shifts are given in ppm (δ) relative to tetramethylsilane as an internal standard. Coupling constants are given in Hz.

#### Synthetic procedures

Scheme S1. Synthesis of BDHI analogs

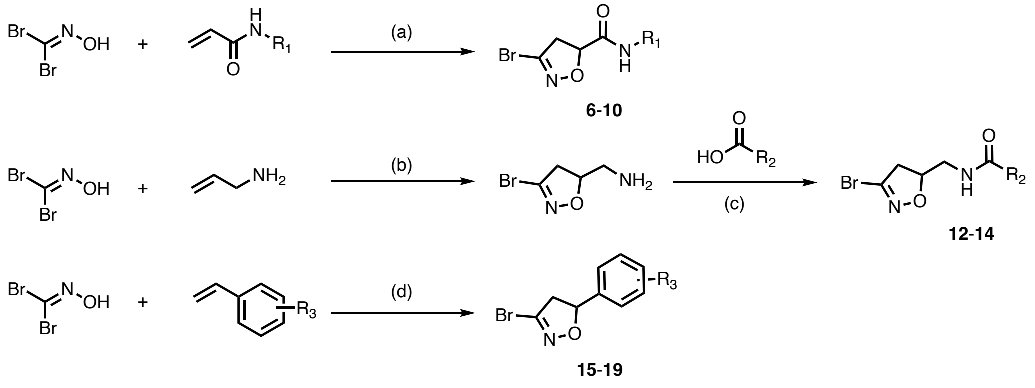


***Reagents and Conditions:*** (a) aq.$KHCO_3$, DMF, rt, 1 h; (b) $KHCO_3$, $H_2O$, rt, 4 h; (c) EDC, DMAP, DMF, rt, 16 h; (d) $KHCO_3$, EtOAc, rt, 16h.

General procedure for synthesis of compounds **6-10** and **24-26**

A solution of acrylamide (0.07 mmol), dibromoformaldoxime (26 mg, 0.13 mmol), and $K_2CO_3$ (68 mg, 0.49 mmol) in EtOAc was stirred at room temperature overnight. The reaction solution was diluted with $H_2O$ and extracted with EtOAc. The crude product was purified by HPLC to give the final product.

*N*-benzyl-3-bromo-4,5-dihydroisoxazole-5-carboxamide (**6**). (28.9 mg, 78%). $^1$H NMR (500 MHz, $CDCl_3$) δ 7.38 – 7.24 (m, 5H), 6.98 (s, 1H), 5.08 (dd, *J* = 11.0, 6.4 Hz, 1H), 4.55 (dd, *J* = 14.7, 6.3 Hz, 1H), 4.38 (dd, *J* = 14.7, 5.4 Hz, 1H), 3.65 – 3.51 (m, 2H). HRMS (ESI): m/z [M + H]$^+$ cald. for $C_{11}H_{12}BrN_2O_2^+$: 283.0077, found: 283.00844.

3-bromo-*N*-phenyl-4,5-dihydroisoxazole-5-carboxamide (**7**). (28.6 mg, 82%). $^1$H NMR (500 MHz, $CDCl_3$) δ 8.37 (s, 1H), 7.57 (d, *J* = 7.5 Hz, 2H), 7.36 (dd, *J* = 8.5, 7.4 Hz, 2H), 7.17 (ddt, *J* = 8.6, 7.3, 1.1 Hz, 1H), 5.17 (dd, *J* = 10.7, 6.6 Hz, 1H), 3.71 – 3.59 (m, 2H). HRMS (ESI): m/z [M + H]$^+$ cald. for $C_{10}H_{10}BrN_2O_2^+$: 268.9920, found: 268.00216.

*N*-(3,5-bis(trifluoromethyl)phenyl)-3-bromo-4,5-dihydroisoxazole-5-carboxamide (**8**). (12.6 mg, 44.4%). $^1$H NMR (500 MHz, $CDCl_3$) δ 8.66 (s, 1H), 8.10 (d, *J* = 1.6 Hz, 2H), 7.67 (s, 1H), 5.20 (dd, *J* = 11.5, 5.5 Hz, 1H), 3.71 (dd, *J* = 18.0, 11.5 Hz, 1H), 3.63 (dd, *J* = 18.0, 5.5 Hz, 1H). HRMS (ESI): m/z [M + H]$^+$ cald. for $C_{12}H_8BrF_6N_2O_2^+$: 404.9668, found: 404.96653.

3-bromo-*N*-(4-methoxyphenyl)-4,5-dihydroisoxazole-5-carboxamide (**9**). (8.6 mg, 26.1%). $^1$H NMR (500 MHz, $CDCl_3$) δ 8.29 (s, 1H), 7.50 – 7.43 (m, 2H), 6.91 – 6.85 (m, 2H), 5.16 (dd, *J* = 10.6, 6.7 Hz, 1H), 3.80 (s, 3H), 3.69 – 3.58 (m, 2H).HRMS (ESI): m/z [M + H]$^+$ cald. for $C_{11}H_{12}BrN_2O_3^+$: 299.0026, found: 299.00615.

(3-bromo-4,5-dihydroisoxazol-5-yl)(6-methoxy-3,4-dihydroquinolin-1(2H)-yl)methanone (**10**). (27.6 mg, 63%). $^1$H NMR (500 MHz, $CDCl_3$) δ 7.28 (s, 1H), 6.81 – 6.70 (m, 2H), 5.37 – 5.26 (m, 1H), 4.17 (dt, *J* = 13.6, 7.1 Hz, 1H), 3.89 – 3.74 (m, 4H), 3.49 (dt, *J* = 13.1, 6.6 Hz, 1H), 3.16 (dd, *J* = 17.0, 10.8 Hz, 1H), 2.73 (dt, *J* = 13.0, 6.2 Hz, 1H), 2.62 (dt, *J* = 14.9, 7.0 Hz, 1H), 2.06 (dt, *J* = 13.0, 6.5 Hz, 1H), 1.87 (dt, *J* = 14.0, 7.2 Hz, 1H). HRMS (ESI): m/z [M + H]$^+$ cald. for $C_{14}H_{16}BrN_2O_3^+$: 339.0339, found: 339.04017.

3-bromo-*N*-(prop-2-yn-1-yl)-4,5-dihydroisoxazole-5-carboxamide (**24**). $^1$H NMR (500 MHz, DMSO) δ 5.07 (dd, *J* = 11.6, 6.9 Hz, 1H), 3.88 (dd, *J* = 5.9, 2.4 Hz, 2H), 3.62 (dd, *J* = 17.6, 11.5 Hz, 1H), 3.37 (dd, *J* =

17.5, 6.9 Hz, 1H), 3.12 (d, *J* = 5.0 Hz, 1H). LC-MS (ESI): m/z $[M + H]^+$ cald. for $C_7H_8BrN_2O_2^+$: 230.9764, found: 230.8.

3-bromo-N-(4-ethynylphenyl)-4,5-dihydroisoxazole-5-carboxamide (**25**). $^1H$ NMR (500 MHz, $CDCl_3$) δ 8.41 (s, 1H), 7.57 – 7.53 (m, 2H), 7.50 – 7.45 (m, 2H), 5.16 (dd, *J* = 11.1, 6.1 Hz, 1H), 3.71 – 3.57 (m, 2H), 3.07 (s, 1H). LC-MS (ESI): m/z $[M + H]^+$ cald. for $C_{12}H_{10}BrN_2O_2^+$: 292.9920, found: 292.8.

(3-bromo-4,5-dihydroisoxazol-5-yl)(6-(hex-5-yn-1-yloxy)-3,4-dihydroquinolin-1(2*H*)-yl)methanone (**26**). (20.42 mg, 29.6%). $^1H$ NMR (500 MHz, $CDCl_3$) δ 6.74 (d, *J* = 10.8 Hz, 2H), 6.11 (s, 1H), 5.36 – 5.28 (m, 1H), 4.16 (dt, *J* = 13.5, 7.0 Hz, 1H), 3.98 (t, *J* = 5.6 Hz, 2H), 3.85 (dd, *J* = 17.0, 8.3 Hz, 1H), 3.50 (dt, *J* = 12.9, 6.5 Hz, 1H), 3.17 (dd, *J* = 16.9, 10.8 Hz, 1H), 2.85 (t, *J* = 7.1 Hz, 2H), 2.76 – 2.69 (m, 1H), 2.66 – 2.58 (m, 1H), 2.06 (dd, *J* = 8.5, 5.3 Hz, 1H), 1.92 – 1.84 (m, 4H), 1.26 (s, 2H). HRMS (ESI): m/z $[M + H]^+$ cald. for $C_{19}H_{22}BrN_2O_3^+$: 405.08083, found: 405.08430.

General procedure for synthesis of compounds **2**-**4** and **12**-**14**

(3-bromo-4,5-dihydroisoxazol-5-yl)methanamine. To a stirred solution of allylamine hydrochloride (230 mg, 2.45 mmol) in water was added dibromoformaldoxime (330 mg, 1.63 mmol). 20% aq. $KHCO_3$ (1 mL) was added dropwise and the solution was stirred for 10 min. 50% aq. KOH was added, and the resulting reaction mixture was filtered and extracted with dichloromethane (DCM, 3 x 10 mL). The combined organic layer was washed with brine, dried over anhydrous $Na_2SO_4$, filtered, and concentrated *in vacuo*. The crude product was purified by CombiFlash (DCM:MeOH, 0 to 10%) to give the title compound as a yellow oil (62 mg, 21%). $^1H$ NMR (500 MHz, $CDCl_3$) δ 4.73 (m, *J* = 10.0, 7.9, 5.8, 3.8 Hz, 1H), 3.24 (dd, *J* = 17.1, 10.6 Hz, 1H), 3.08 (dd, *J* = 17.1, 8.0 Hz, 1H), 3.00 (dd, *J* = 13.6, 3.8 Hz, 1H), 2.85 (dd, *J* = 13.6, 5.8 Hz, 1H). LC-MS (ESI): m/z $[M + H]^+$ calcd. for $C_4H_8BrN_2O^+$: 178.98, found: 179.1.

A solution of carboxylic acid (0.06 mmol), (3-bromo-4,5-dihydroisoxazol-5-yl)methanamine (11 mg, 0.064 mmol), EDC (14 mg, 0.09 mmol), and DMAP (11 mg, 0.09 mmol) in DMF was stirred at room temperature overnight. DMF was evaporated, and the crude reaction mixture was purified by HPLC to give the title compound.

5-(((3-bromo-4,5-dihydroisoxazol-5-yl)methyl)carbamoyl)-2-(6-(dimethylamino)-3-(dimethyliminio)-3H-xanthen-9-yl)benzoate (**2**). (1.87 mg, 83%). LC-MS (ESI): m/z $[M + H]^+$ calcd. for $C_{29}H_{28}BrN_4O_5^+$: 591.1, found: 590.8.

5-((1-(3-bromo-4,5-dihydroisoxazol-5-yl)-3-oxo-6,9,12,15-tetraoxa-2-azaheptadecan-17-yl)carbamoyl)-2-(6-(dimethylamino)-3-(dimethyliminio)-3H-xanthen-9-yl)benzoate (**3**). (0.67 mg, 54%). LC-MS (ESI): m/z $[M + H]^+$ calcd. for $C_{40}H_{49}BrN_5O_{10}^+$: 838.3, found: 837.8.

*N*-((3-bromo-4,5-dihydroisoxazol-5-yl)methyl)pent-4-ynamide (**4**). White solid (4.25 mg, 54%). $^1H$ NMR (500 MHz, $CDCl_3$) δ 5.95 (m, 1H), 4.86 – 4.77 (m, 1H), 3.68 – 3.52 (m, 2H), 3.27 (dd, *J* = 17.5, 10.7 Hz, 1H), 3.02 (dd, *J* = 17.5, 8.1 Hz, 1H), 2.54 (ddt, *J* = 10.2, 6.2, 2.8 Hz, 2H), 2.48 – 2.38 (m, 2H), 2.03 (t, *J* = 2.6 Hz, 1H). HRMS (ESI): m/z $[M + H]^+$ cald. for $C_9H_{12}BrN_2O_2^+$: 259.0077, found: 259.00767.

*N*-((3-bromo-4,5-dihydroisoxazol-5-yl)methyl)-3,5-bis(trifluoromethyl)benzamide (**12**). (5 mg, 20.6%). $^1H$ NMR (500 MHz, $CDCl_3$) δ 8.23 – 8.21 (m, 2H), 8.04 (d, *J* = 1.8 Hz, 1H), 6.64 (s, 1H), 4.94 (dtd, *J* = 10.3, 6.8, 3.1 Hz, 1H), 3.85 (dd, *J* = 14.4, 6.2, 3.1 Hz, 1H), 3.71 (dd, *J* = 14.4, 6.3 Hz, 1H), 3.41 (dd, *J* = 17.5, 10.6 Hz, 1H), 3.06 (dd, *J* = 17.6, 7.1 Hz, 1H). HRMS (ESI): m/z $[M + H]^+$ cald. for $C_{13}H_{10}BrF_6N_2O_2^+$: 418.9824, found: 418.98816.

*N*-((3-bromo-4,5-dihydroisoxazol-5-yl)methyl)-2-(pyridin-3-yl)-4,5-dihydrothiazole-4-carboxamide (**13**). $^1H$ NMR (500 MHz, $CDCl_3$) δ 9.15 (t, *J* = 3.0 Hz, 1H), 8.87 – 8.77 (m, 1H), 8.53 – 8.41 (m, 1H), 7.72 (ddt, *J* = 13.8, 8.1, 5.4 Hz, 1H), 7.11 (d, *J* = 43.9 Hz, 2H), 5.32 – 5.22 (m, 1H), 4.99 – 4.74 (m, 2H), 3.89 – 3.50 (m, 5H), 3.43 – 3.26 (m, 2H), 3.07 (dddd, *J* = 28.4, 17.5, 7.6, 3.3 Hz, 1H), 2.94 (ddd, *J* = 17.6, 7.0, 3.2 Hz, 1H). HRMS (ESI): m/z $[M + H]^+$ cald. for $C_{13}H_{14}BrN_4O_2S^+$: 369.0015, found: 369.01351.

N-((3-bromo-4,5-dihydroisoxazol-5-yl)methyl)-1,3-dimethyl-1H-pyrazolo[3,4-b]pyridine-5-carboxamide (**14**). $^{1}H$ NMR (500 MHz, $CDCl_3$) δ 8.85 (d, *J* = 2.1 Hz, 1H), 8.35 (d, *J* = 2.1 Hz, 1H), 6.51 – 6.45 (m, 1H), 4.88 (m, *J* = 10.6, 7.4, 5.8, 3.3 Hz, 1H), 4.03 (s, 3H), 3.76 (ddd, *J* = 14.5, 6.0, 3.3 Hz, 1H), 3.70 (dt, *J* = 14.4, 6.1 Hz, 1H), 3.31 (dd, *J* = 17.5, 10.7 Hz, 1H), 3.02 (dd, *J* = 17.5, 7.3 Hz, 1H), 2.52 (s, 3H). HRMS (ESI): m/z $[M + H]^+$ cald. for $C_{13}H_{15}BrN_5O_2^+$: 352.0404, found: 352.0485.

General procedure for synthesis of compounds **5** and **15**-**19**
A solution of styrene (0.29 mmol), dibromoformaldoxime (101 mg, 0.52 mmol), and $K_2CO_3$ (281 mg, 2.0 mmol) in EtOAc was stirred at room temperature overnight. The reaction solution was the diluted with $H_2O$ and extracted with EtOAc. The crude product was purified by CombiFlash (Hx:EtOAc, 0 to 20%) to give the final product.

3-(3-bromo-4,5-dihydroisoxazol-5-yl)-*N*-(prop-2-yn-1-yl)benzamide (**5**). (9.3 mg, 13.8%). $^{1}H$ NMR (500 MHz, $CDCl_3$) δ 7.75 (dt, *J* = 8.8, 1.9 Hz, 2H), 7.53 – 7.45 (m, 2H), 6.46 (t, *J* = 5.4 Hz, 1H), 5.73 (dd, *J* = 10.9, 8.8 Hz, 1H), 4.26 (dd, *J* = 5.3, 2.5 Hz, 2H), 3.66 (dd, *J* = 17.3, 11.0 Hz, 1H), 3.20 (dd, *J* = 17.3, 8.8 Hz, 1H), 2.30 (d, *J* = 2.4 Hz, 1H). HRMS (ESI): m/z $[M + H]^+$ cald. for $C_{13}H_{12}BrN_2O_2^+$: 307.00767, found: 307.01105.

3-bromo-5-phenyl-4,5-dihydroisoxazole (**15**). (35.3 mg, 53.8%). $^{1}H$ NMR (500 MHz, $CDCl_3$) δ 7.43 – 7.32 (m, 5H), 5.67 (dd, *J* = 10.9, 9.0 Hz, 1H), 3.62 (dd, *J* = 17.2, 10.9 Hz, 1H), 3.21 (dd, *J* = 17.2, 9.0 Hz, 1H). HRMS (ESI): m/z $[M + H]^+$ cald. for $C_9H_9BrNO^+$: 225.9862, found: 225.98654.

3-bromo-5-(3-nitrophenyl)-4,5-dihydroisoxazole (**16**). (6.2 mg, 17.6%). $^{1}H$ NMR (500 MHz, $CDCl_3$) δ 8.25 – 8.19 (m, 2H), 7.74 – 7.68 (m, 1H), 7.64 – 7.57 (m, 1H), 5.78 (dd, *J* = 11.0, 8.4 Hz, 1H), 3.74 (dd, *J* = 17.3, 11.0 Hz, 1H), 3.21 (dd, *J* = 17.3, 8.4 Hz, 1H). HRMS (ESI): m/z $[M + H]^+$ cald. for $C_9H_8BrN_2O_3^+$: 270.9713, found: 270.97153.

3-bromo-5-(4-nitrophenyl)-4,5-dihydroisoxazole (**17**). (18.6 mg, 52.8%). $^{1}H$ NMR (500 MHz, $CDCl_3$) δ 8.29 – 8.22 (m, 2H), 7.56 – 7.49 (m, 2H), 5.78 (dd, *J* = 11.1, 8.2 Hz, 1H), 3.74 (dd, *J* = 17.2, 11.1 Hz, 1H), 3.17 (dd, *J* = 17.3, 8.2 Hz, 1H). HRMS (ESI): m/z $[M + H]^+$ cald. for $C_9H_8BrN_2O_3^+$: 270.9713, found: 270.97120.

3-bromo-5-(4-methoxyphenyl)-4,5-dihydroisoxazole (**18**). (9.7 mg, 25.3%). $^{1}H$ NMR (500 MHz, $CDCl_3$) δ 7.29 – 7.26 (m, 2H), 6.95 – 6.87 (m, 2H), 5.62 (dd, *J* = 10.8, 9.4 Hz, 1H), 3.82 (s, 3H), 3.56 (dd, *J* = 17.2, 10.8 Hz, 1H), 3.20 (dd, *J* = 17.3, 9.4 Hz, 1H). HRMS (ESI): m/z $[M + H]^+$ cald. for $C_{10}H_{11}BrNO_2^+$: 255.9968, found: 255.99728.

5-(3,5-bis(trifluoromethyl)phenyl)-3-bromo-4,5-dihydroisoxazole (**19**). (2.9 mg, 20%). $^{1}H$ NMR (500 MHz, $CDCl_3$) δ 7.88 (s, 1H), 7.81 (s, 2H), 5.80 (dd, *J* = 11.1, 8.3 Hz, 1H), 3.76 (dd, *J* = 17.3, 11.1 Hz, 1H), 3.21 (dd, *J* = 17.3, 8.3 Hz, 1H). HRMS (ESI): m/z $[M + H]^+$ cald. for $C_{11}H_7BrF_6NO^+$: 361.9610, found: 361.96117.

*N*-(2-(2-iodoacetamido)ethyl)-6-(5-methyl-2-oxoimidazolidin-4-yl)hexanamide (IA-DTB). IA-DTB was prepared following a reported procedure. $^{1}H$ NMR (500 MHz, DMSO) δ 8.26 (s, 1H), 7.78 (s, 1H), 6.29 (s, 1H), 6.11 (s, 1H), 3.62 – 3.56 (m, 3H), 3.50 – 3.44 (m, 1H), 3.08 (q, *J* = 2.7, 2.2 Hz, 4H), 2.04 (t, *J* = 7.5 Hz, 2H), 1.47 (q, *J* = 7.4 Hz, 2H), 1.37 – 1.28 (m, 3H), 1.28 – 1.16 (m, 3H), 0.95 (d, *J* = 6.5 Hz, 3H). LC-MS (ESI): m/z $[M + H]^+$ calcd. for $C_{14}H_{26}IN_4O_3^+$: 425.1, found: 425.2.

### NMR data

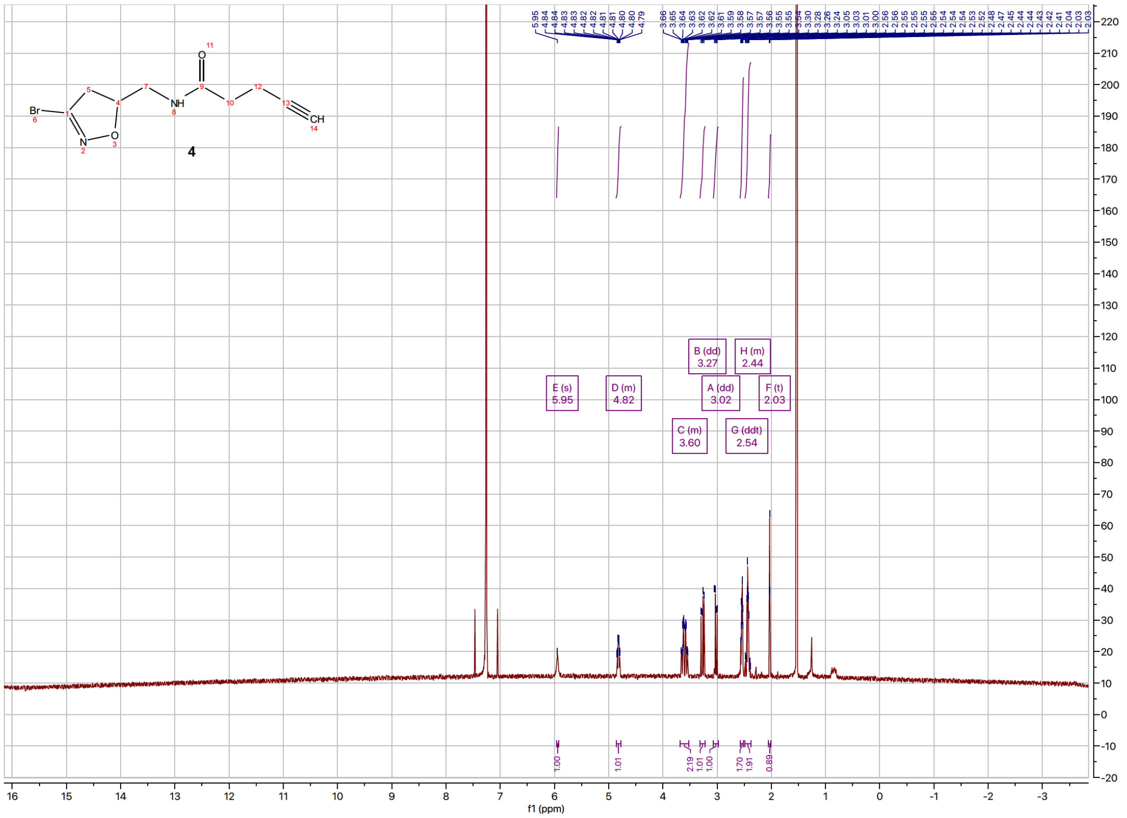
4
E (s) 5.95
D (m) 4.82
C (m) 3.60
B (dd) 3.27
A (dd) 3.02
G (ddt) 2.54
H (m) 2.44
F (t) 2.03
f1 (ppm)

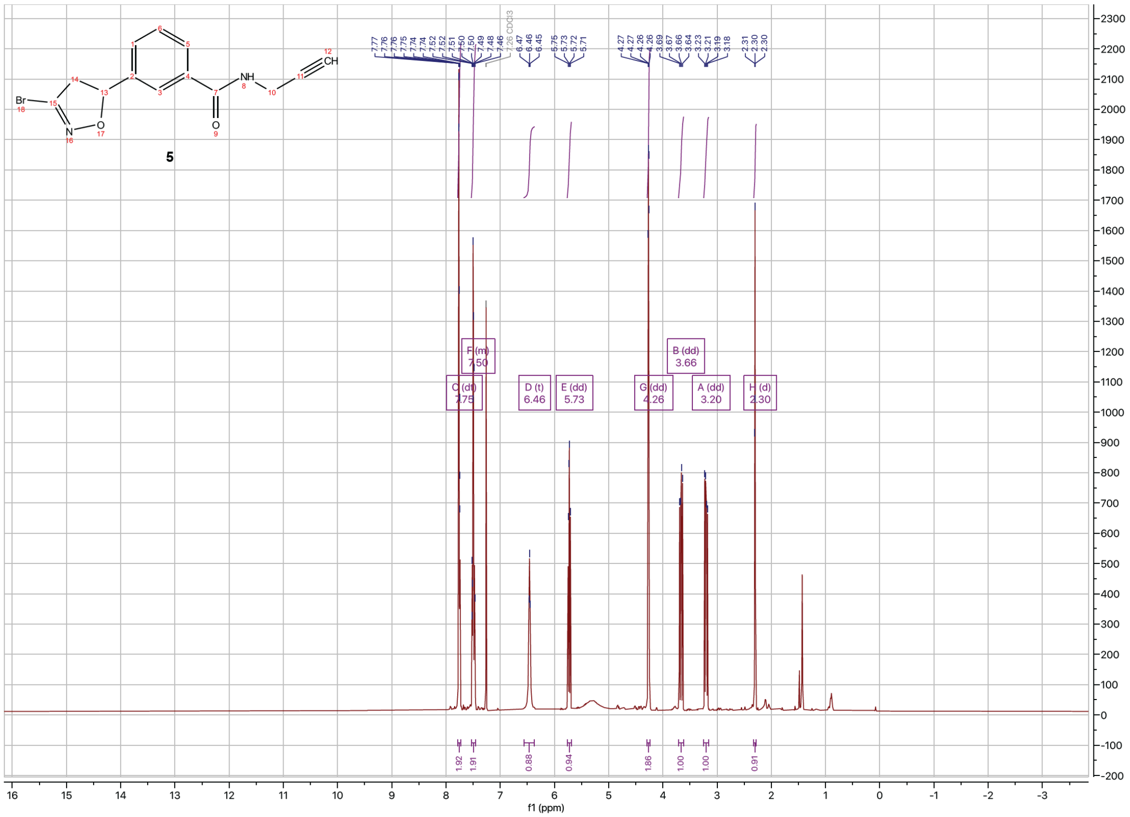
5
C (dt) 7.75
F (m) 7.50
D (t) 6.46
E (dd) 5.73
G (dd) 4.26
B (dd) 3.66
A (dd) 3.20
H (d) 2.30
f1 (ppm)

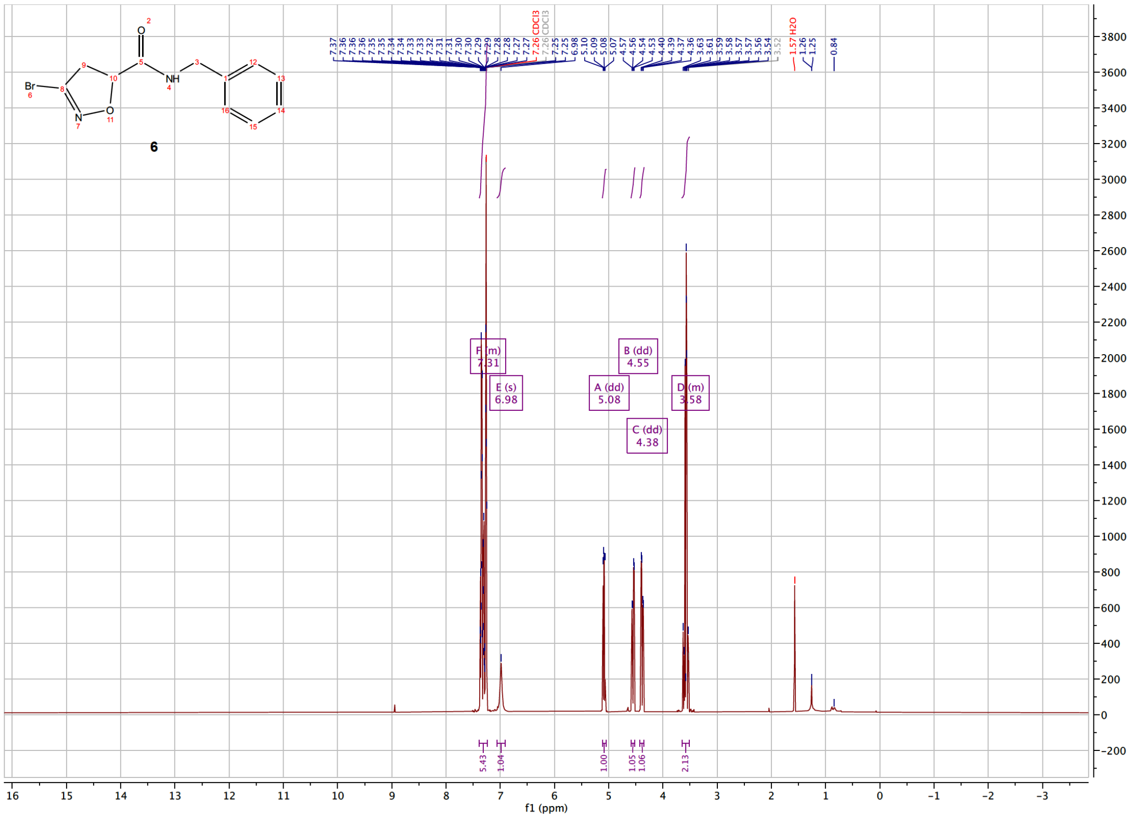

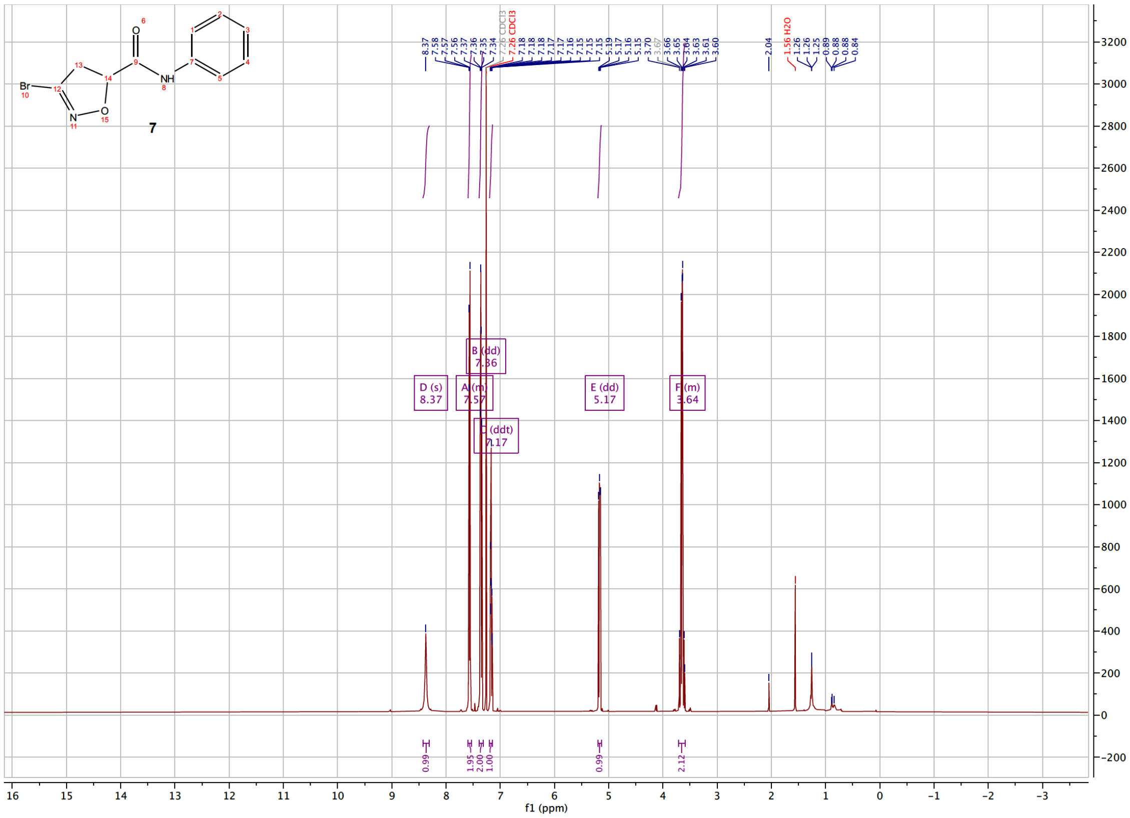

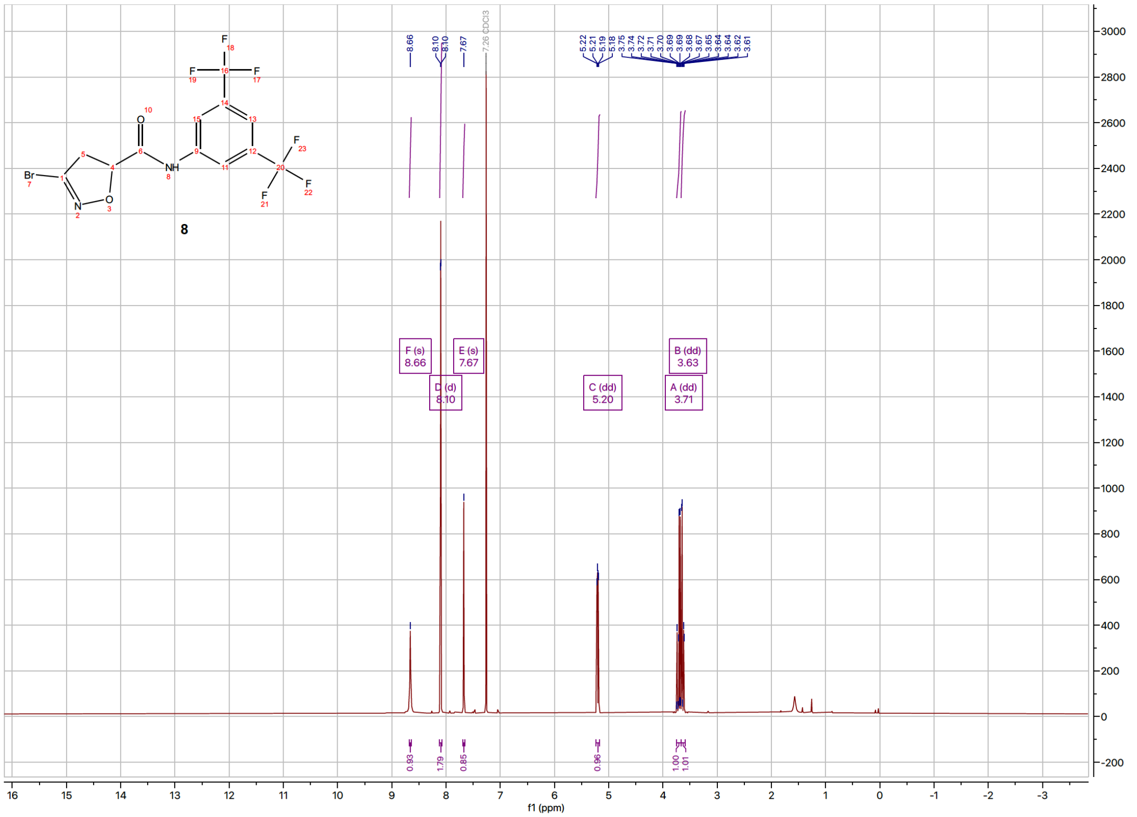
8
F (s)
8.66
D (d)
8.10
E (s)
7.67
C (dd)
5.20
B (dd)
3.63
A (dd)
3.71
f1 (ppm)

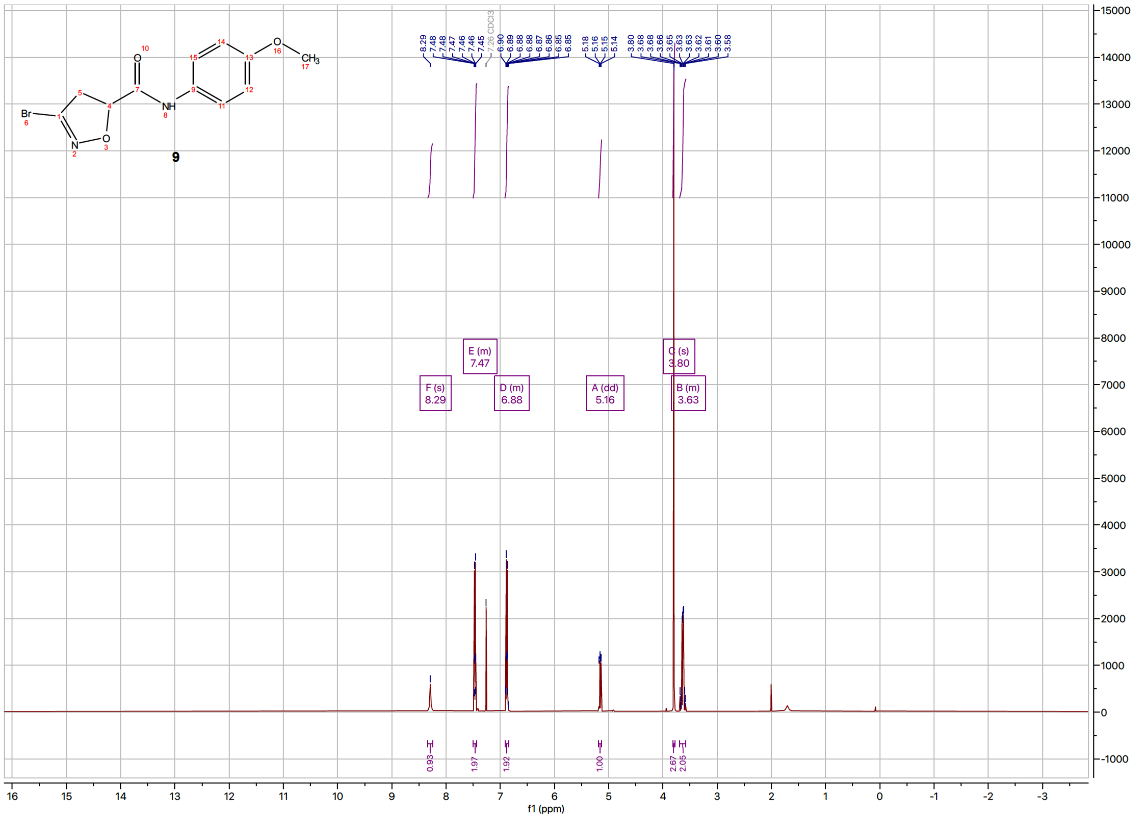
9
F (s)
8.29
E (m)
7.47
D (m)
6.88
A (dd)
5.16
C (s)
3.80
B (m)
3.63
f1 (ppm)

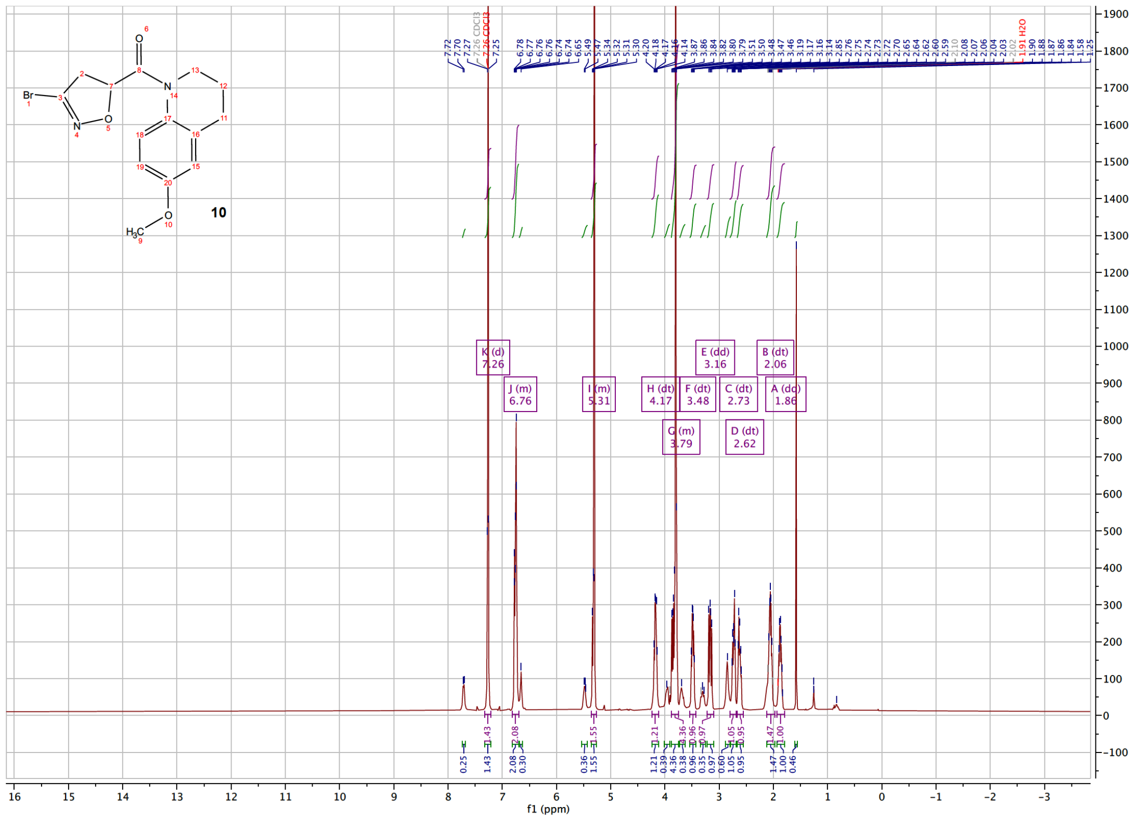
10
K (d)
7.26
J (m)
6.76
I (m)
5.31
H (dt)
4.17
G (m)
3.79
F (dt)
3.48
E (dd)
3.16
C (dt)
2.73
D (dt)
2.62
B (dt)
2.06
A (dd)
1.86
f1 (ppm)

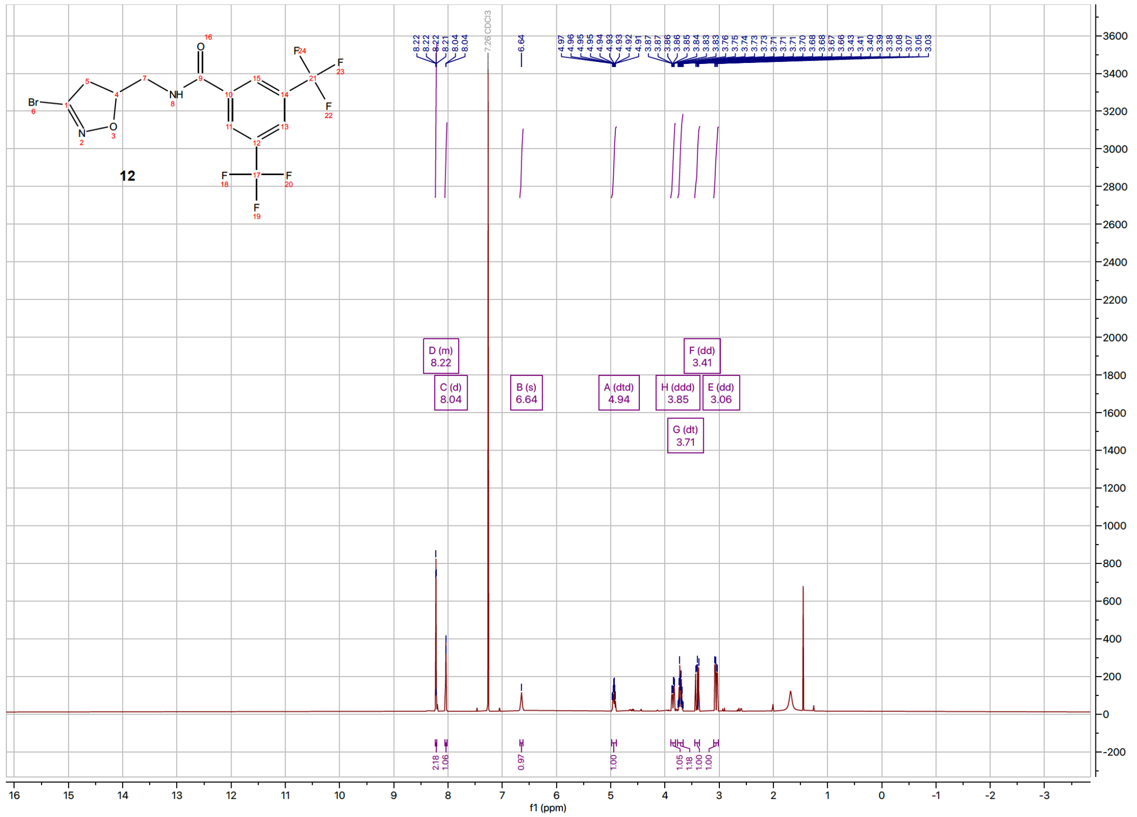
12
D (m)
8.22
C (d)
8.04
B (s)
6.64
A (dtd)
4.94
H (ddd)
3.85
G (dt)
3.71
F (dd)
3.41
E (dd)
3.06
f1 (ppm)

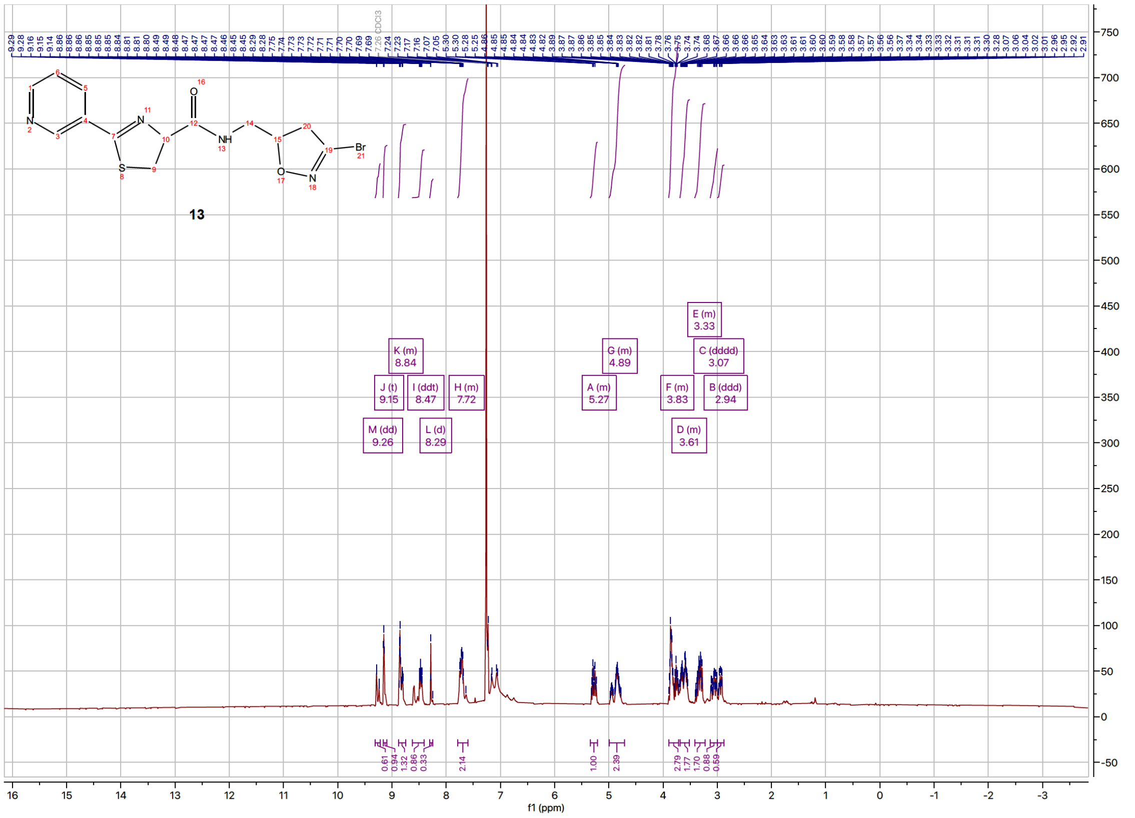

13
CDCl3
K (m) 8.84
G (m) 4.89
E (m) 3.33
C (dddd) 3.07
J (t) 9.15
I (ddt) 8.47
H (m) 7.72
A (m) 5.27
F (m) 3.83
B (ddd) 2.94
M (dd) 9.26
L (d) 8.29
D (m) 3.61
f1 (ppm)


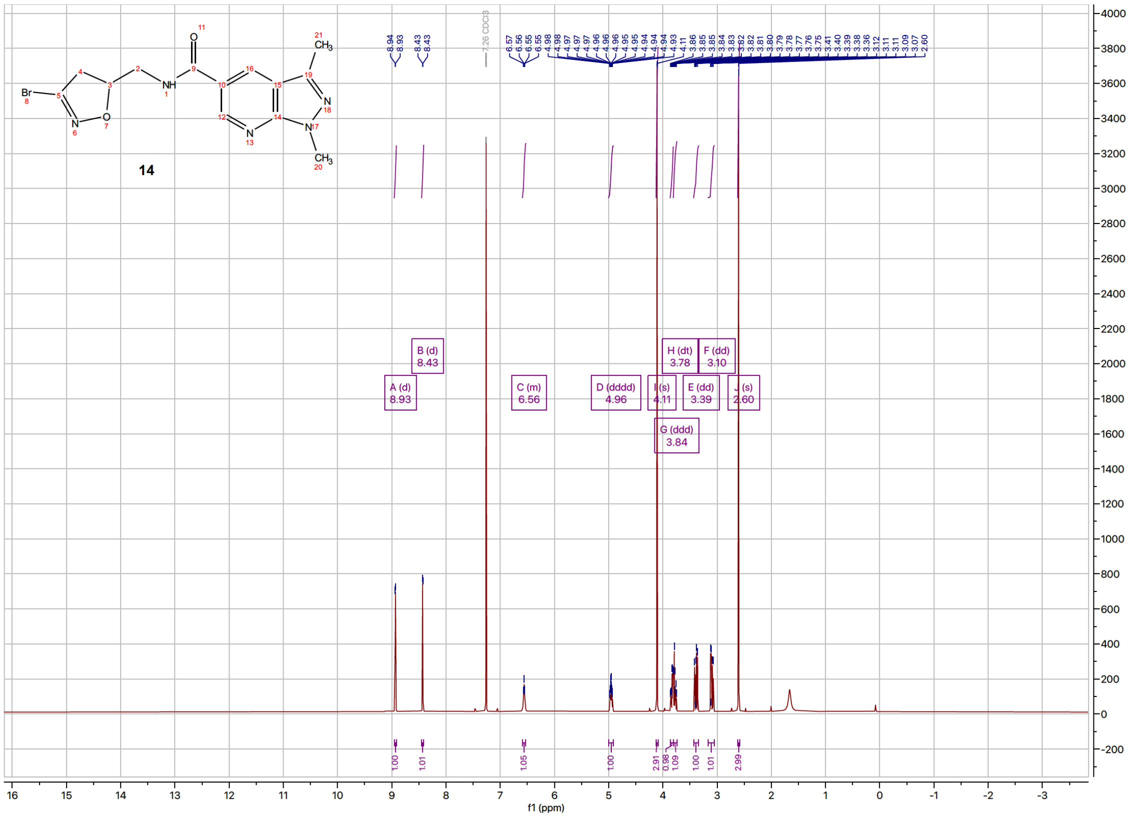

14
7.26 CDCl3
B (d) 8.43
A (d) 8.93
C (m) 6.56
D (dddd) 4.96
I (s) 4.11
H (dt) 3.78
F (dd) 3.10
E (dd) 3.39
J (s) 2.60
G (ddd) 3.84
f1 (ppm)

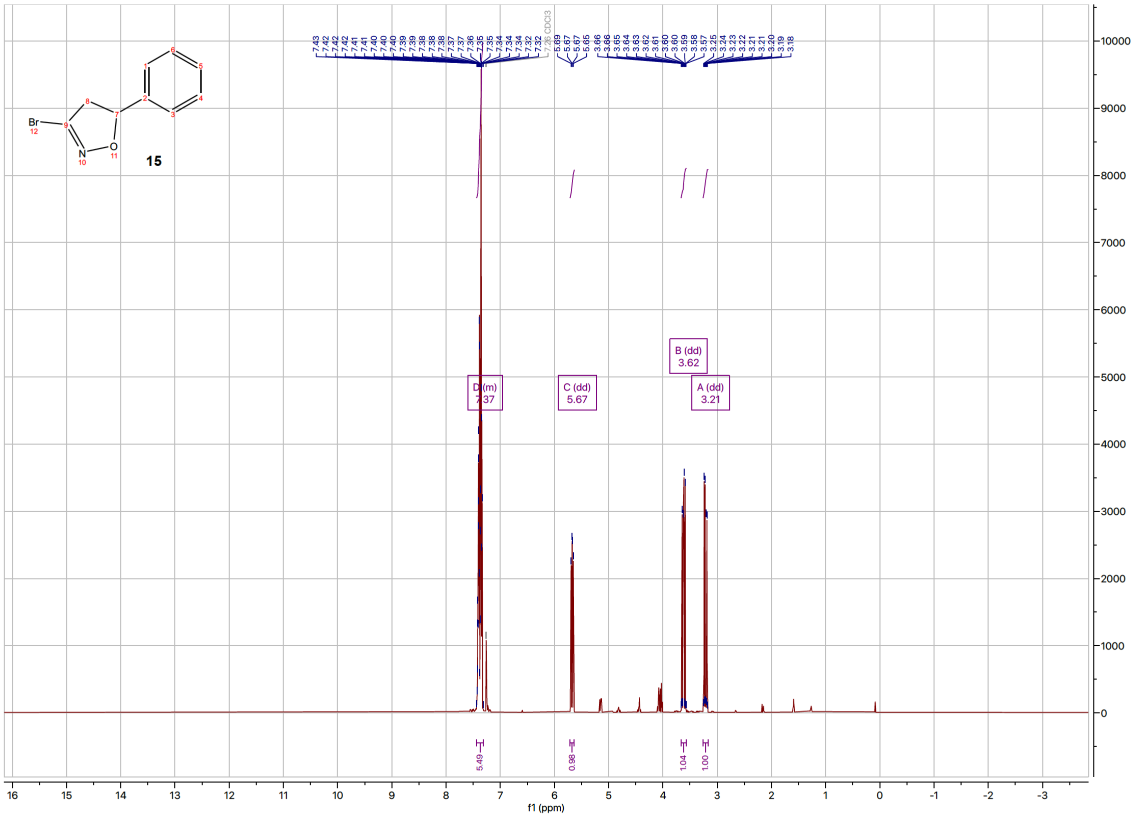
15
Br
N
O
7.43 7.42 7.42 7.41 7.41 7.40 7.40 7.40 7.39 7.39 7.38 7.38 7.38 7.37 7.37 7.36 7.35 7.35 7.34 7.34 7.34 7.32 7.32
7.26 CDCl3
5.69 5.67 5.67 5.65
3.66 3.66 3.65 3.64 3.63 3.62 3.61 3.60 3.60 3.59 3.58 3.57 3.25 3.24 3.23 3.22 3.21 3.21 3.20 3.19 3.18
D (m) 7.37
C (dd) 5.67
B (dd) 3.62
A (dd) 3.21
5.49
0.98
1.04
1.00
f1 (ppm)

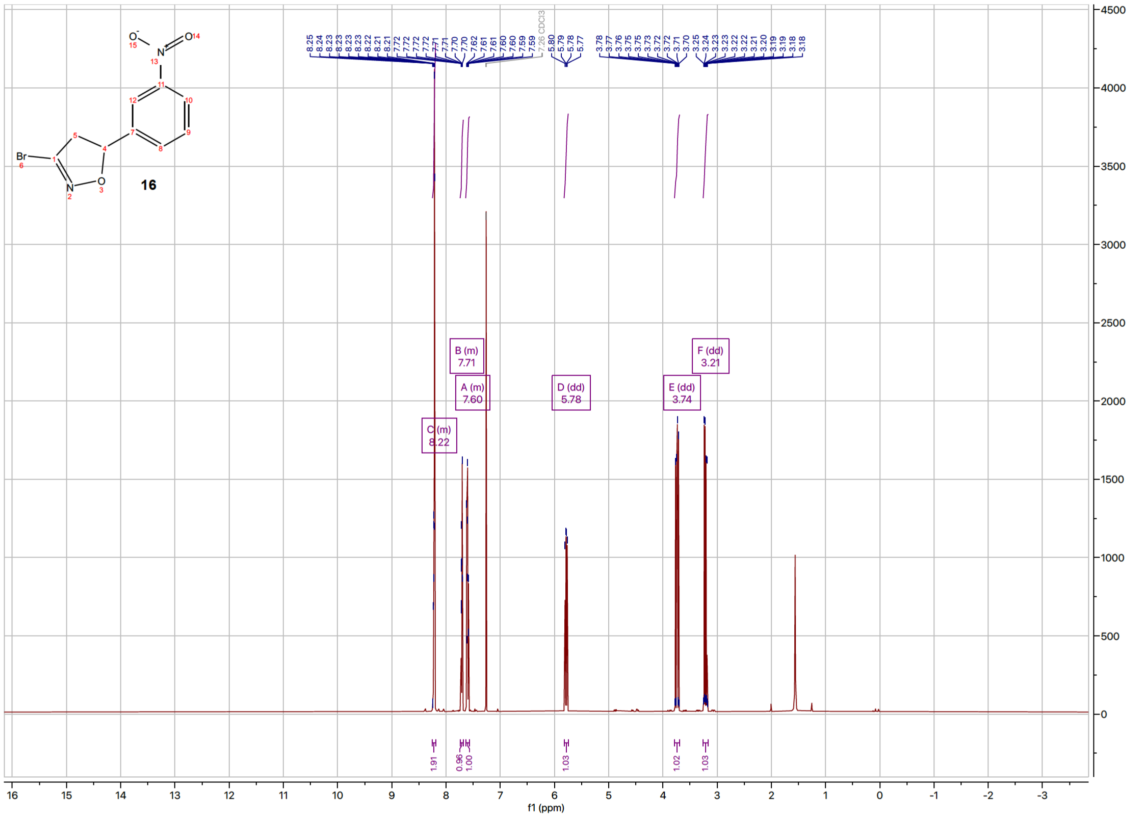
16
Br
N
O
8.25 8.24 8.23 8.23 8.23 8.22 8.21 8.21 7.72 7.72 7.72 7.71 7.71 7.70 7.70 7.62 7.61 7.61 7.60 7.60 7.59 7.59
7.26 CDCl3
5.80 5.79 5.78 5.77
3.78 3.77 3.76 3.75 3.75 3.73 3.72 3.72 3.71 3.70 3.25 3.24 3.23 3.23 3.22 3.22 3.21 3.20 3.19 3.19 3.18 3.18
C (m) 8.22
B (m) 7.71
A (m) 7.60
D (dd) 5.78
E (dd) 3.74
F (dd) 3.21
1.91
0.96
1.00
1.03
1.02
1.03
f1 (ppm)

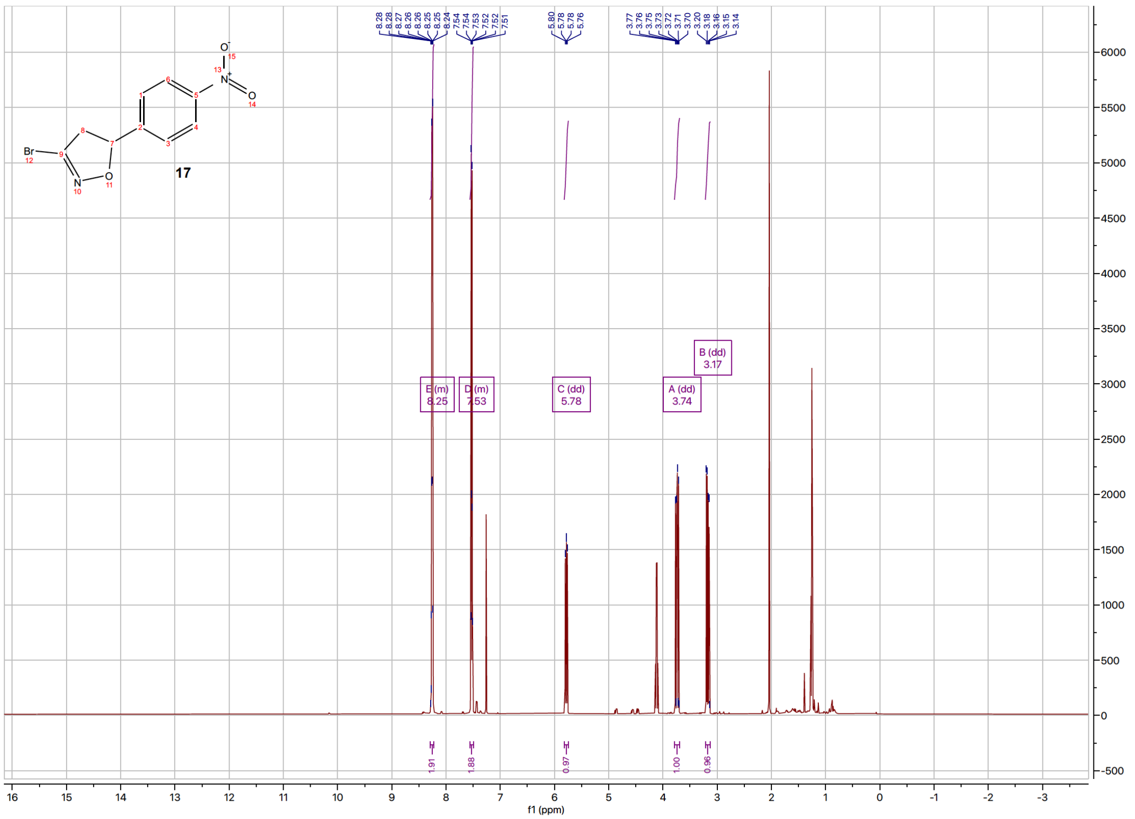

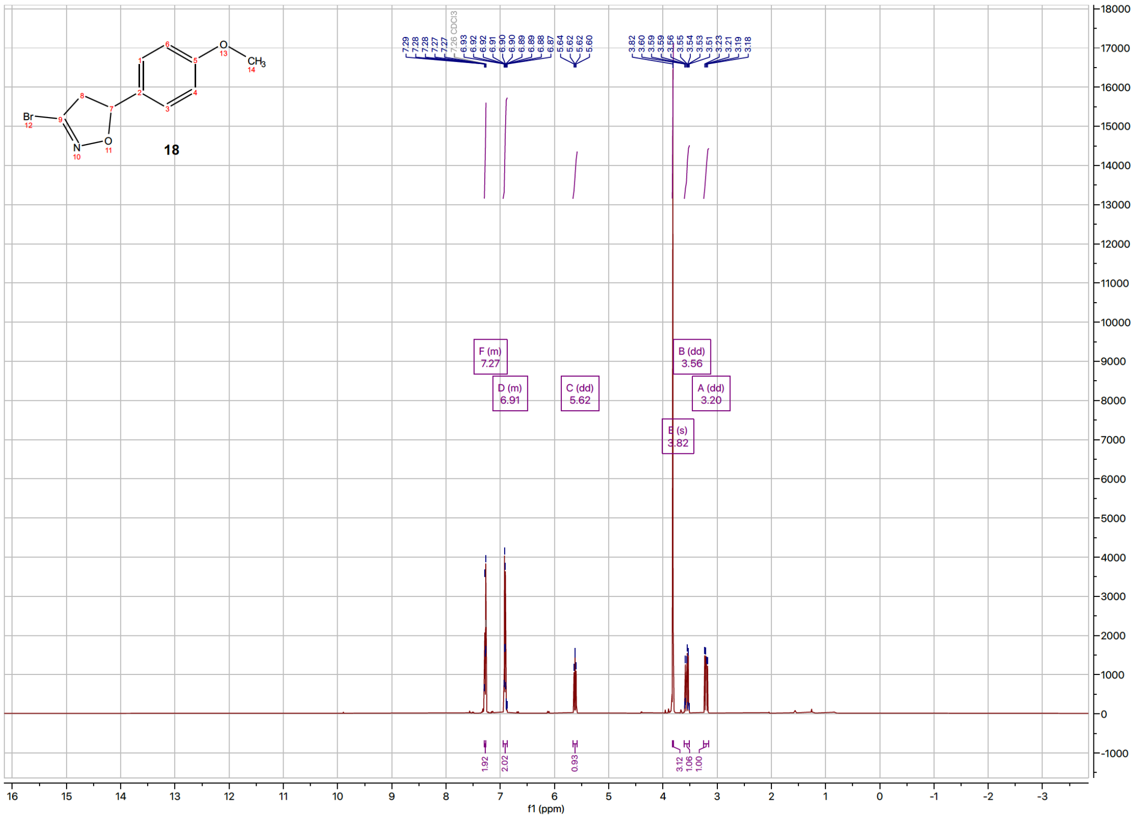

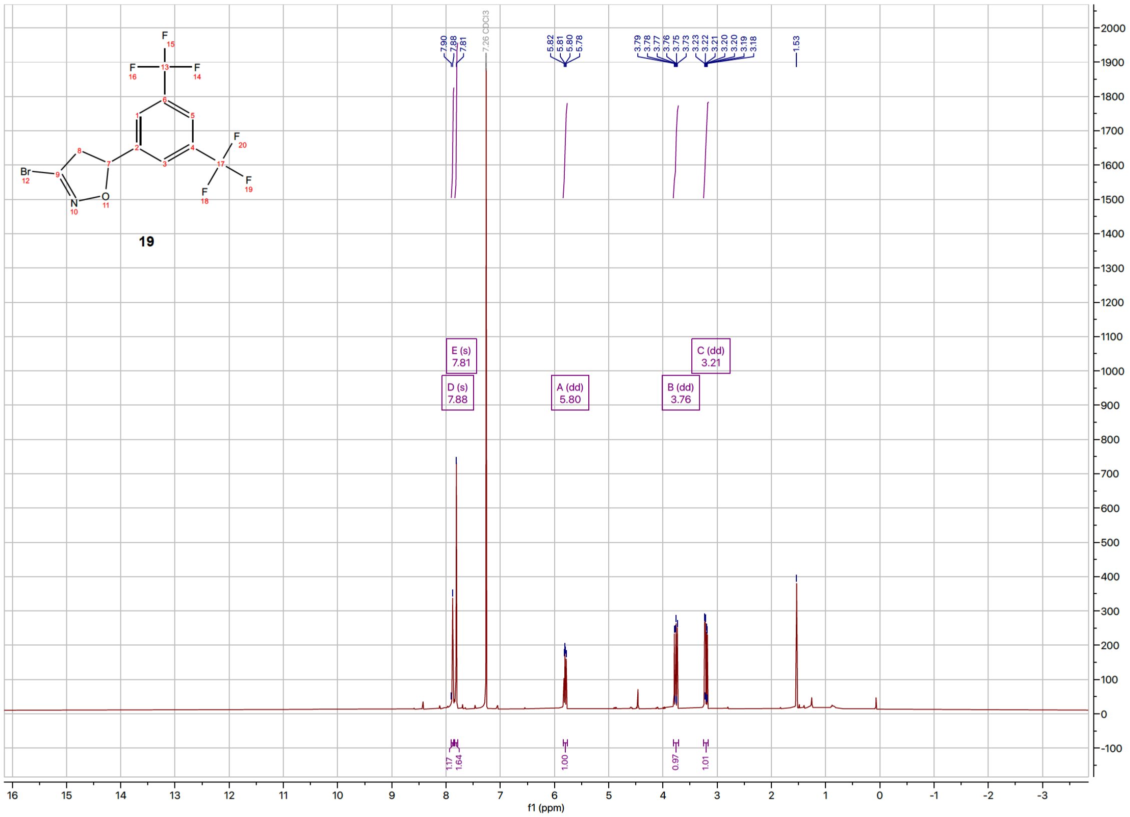
19
7.90
7.88
7.81
7.26 CDCl3
5.82
5.81
5.80
5.78
3.79
3.78
3.77
3.76
3.75
3.73
3.23
3.22
3.21
3.20
3.19
3.18
1.53
E (s) 7.81
D (s) 7.88
A (dd) 5.80
B (dd) 3.76
C (dd) 3.21
1.17
1.64
1.00
0.97
1.01
f1 (ppm)

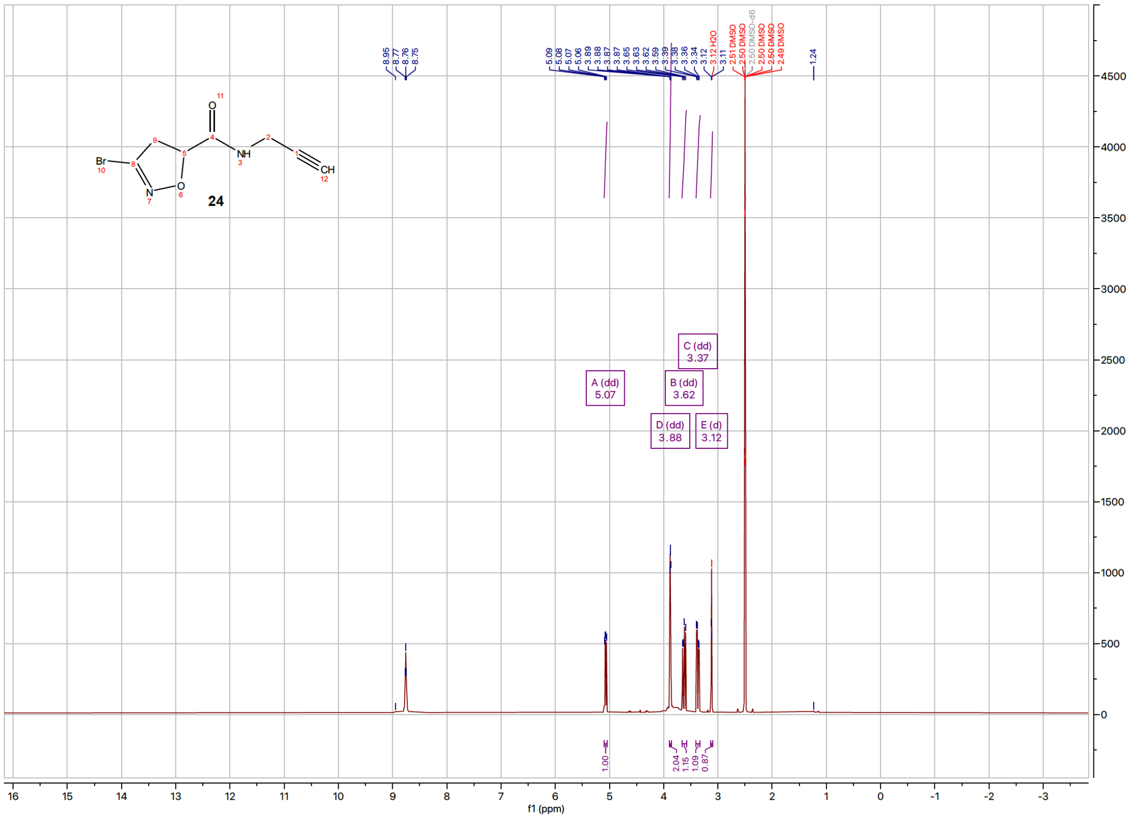
24
8.95
8.77
8.76
8.75
5.09
5.08
5.07
5.06
3.89
3.88
3.87
3.87
3.65
3.63
3.62
3.59
3.39
3.38
3.36
3.34
3.12
3.11
3.12 H2O
2.51 DMSO
2.50 DMSO
2.50 DMSO-d6
2.50 DMSO
2.50 DMSO
2.49 DMSO
1.24
A (dd) 5.07
B (dd) 3.62
C (dd) 3.37
D (dd) 3.88
E (d) 3.12
1.00
2.04
1.15
1.09
0.87
f1 (ppm)

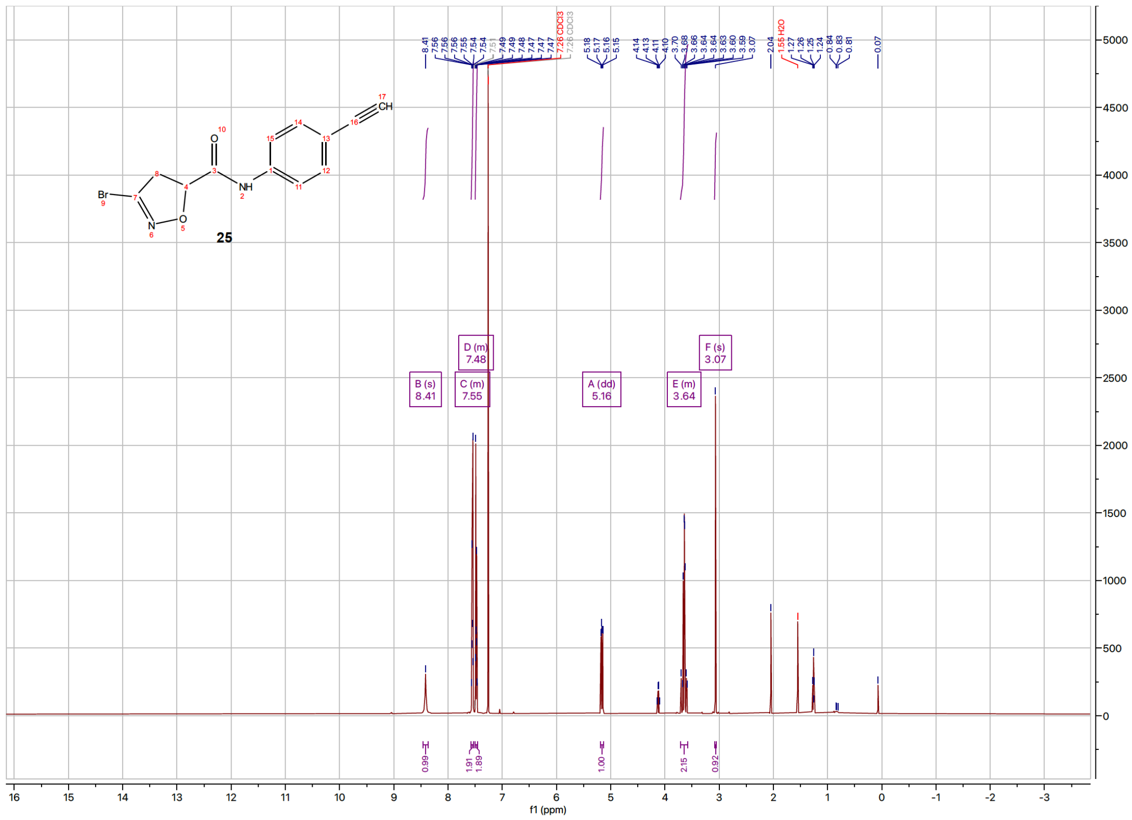

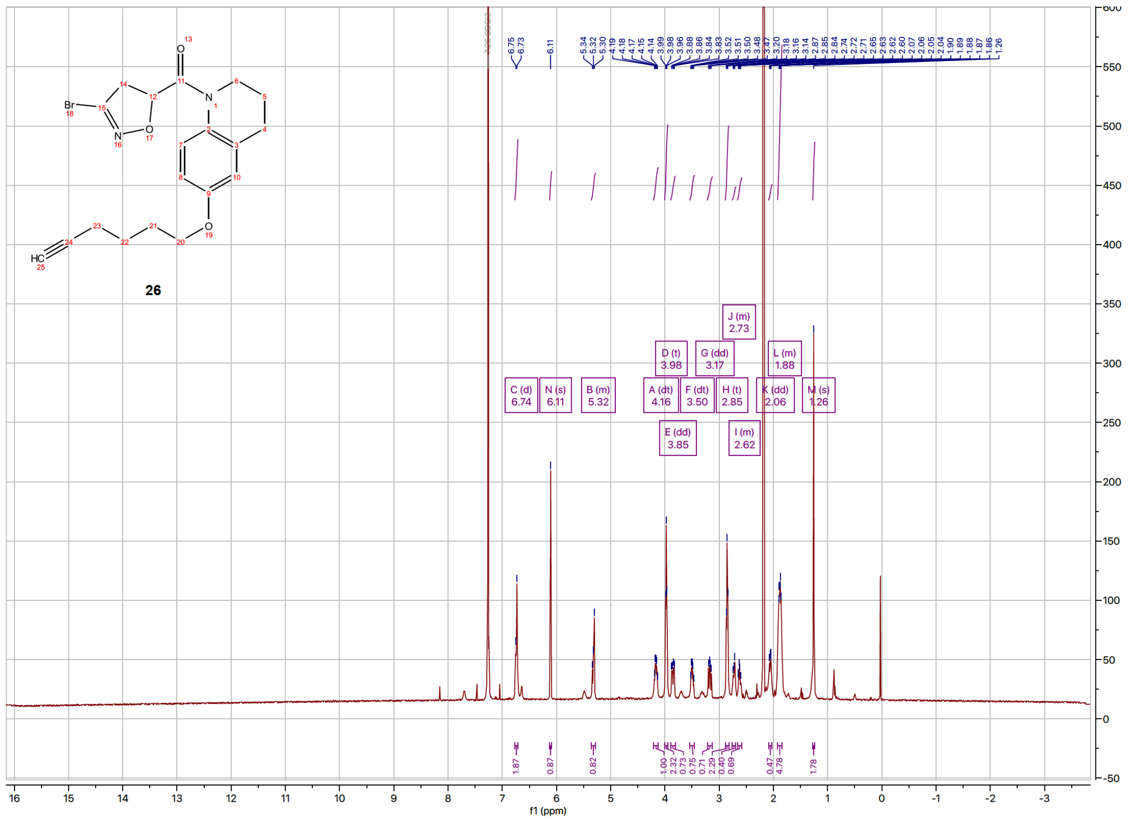

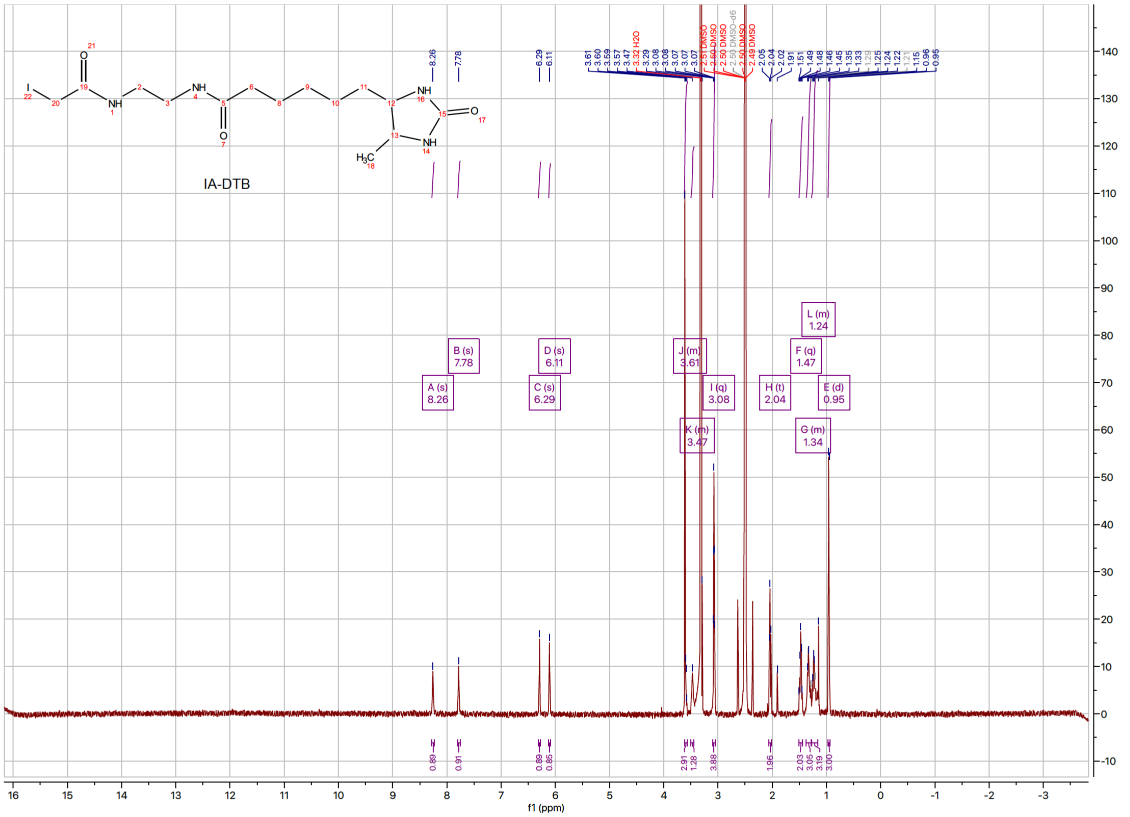
IA-DTB
A (s)
8.26
B (s)
7.78
C (s)
6.29
D (s)
6.11
J (m)
3.61
K (m)
3.47
I (q)
3.08
H (t)
2.04
F (q)
1.47
L (m)
1.24
G (m)
1.34
E (d)
0.95
f1 (ppm)
